# Mechanism of Lipid-Dependent Thermosensation in an Ion Channel

**DOI:** 10.1101/2025.06.03.657524

**Authors:** Chieh-Chin Li, Crina M. Nimigean

## Abstract

Temperature sensing is a fundamental biological process that allows organisms to detect and respond to environmental. Ion channels that gate in response to temperature changes, known as thermoreceptors, play key roles in this process. Although the phenomena affected by temperature in many thermoreceptors have been widely studied, the molecular mechanisms by which these proteins sense temperature remain controversial and poorly understood. Here, we investigated the molecular details of temperature sensing in a thermophilic bacterial ion channel, SthK from *Spirochaeta thermophila*, a homologue of mammalian hyperpolarization-activated and cyclic nucleotide-modulated (HCN) channels with well-characterized structure and function. We show that SthK is a cold-sensitive ion channel, displaying higher activity when temperatures are decreased below 30 °C. Intriguingly, the SthK cold sensitivity is highly dependent on the lipid composition, being sensitive in the presence of amine-containing lipids, and insensitive in the presence of anionic lipids. By combining cryo-EM structural analysis, mutagenesis, and functional assays, we demonstrated that a functionally important intersubunit salt bridge that was previously shown to be important in lipid modulation, acts as the temperature sensor in these channels. This salt bridge is state-dependent, stabilizes closed states, and needs to break for the channel to open. Lower temperatures that weaken salt bridge interactions thus favor channel opening. Lipid-binding at this location tune temperature sensitivity by modulating the salt bridge strength through their headgroup charge and size. Interesting to note, equivalent salt bridges are also found in other ion channels such as HCN channels and, notably, TRPM8, an established cold-sensitive thermoreceptor, suggesting that this mechanism could be conserved in other temperature-sensitive ion channels. Our findings highlight a finely tuned interplay between structural elements and membrane environment, suggesting that thermosensitivity can emerge from the cooperative effects of salt bridge energetics and lipid context.

## Introduction

Temperature sensing is essential for organismal survival, enabling the detection of environmental fluctuations and the maintenance of physiological homeostasis [1]. In mammals, thermal stimuli are primarily detected by somatosensory neurons that express temperature-gated ion channels. Among these, the transient receptor potential (TRP) channel family, especially the subset known as thermo-TRPs, has been the dominant focus of thermosensation research due to their striking temperature-dependent gating properties and clear physiological relevance in heat and cold sensing [2]. These channels include both heat- and cold-sensitive members; for example, TRPV1 responds to noxious heat, TRPV3 and TRPV4 to innocuous warmth, and TRPM8 and TRPA1 to cold, among others, forming a modular system for detecting a wide range of thermal cues.

Despite their shared ability to detect thermal stimuli, the molecular basis of temperature gating in TRP channels remains elusive. Multiple regions have been implicated in thermosensitivity, such as the pore domain, pore turret, N-terminal ankyrin repeats, and C-terminal tail, although experimental findings are often conflicting. For example, the pore turret has been proposed to contribute to TRPV1 heat activation, yet other studies suggest it is dispensable [3, 4]. Similarly, the C-terminal domain appears to influence temperature tuning in TRPV1–TRPM8 chimeras, but heat sensitivity persists in vampire bat TRPV1 variants lacking most of this region [5, 6]. These inconsistencies raised the possibility that no single conserved domain underlies thermosensation. In that vein, Clapham and Miller proposed that temperature gating arises from a large difference in heat capacity (Δ*Cₚ*) between channel states, contributed by multiple elements within the channels rather than from a discrete temperature sensor [7]. This model is supported by a rationally-designed Shaker channel study, which demonstrated that temperature sensitivity can arise from changes in heat capacity from strategically-introduced residues rather than from a specialized domain [8].

Structurally, some of the thermosensitive channels have been shown to respond to temperature by undergoing conformational rearrangements [9, 10] and others by employing temperature-dependent ligand binding [11], although the molecular mechanism is not clear. Beyond thermo-TRPs, temperature-dependent reversible unfolding of a cytoplasmic domain has also been shown to control voltage-dependent gating in a bacterial voltage-gated sodium channel [12], highlighting an alternative structural mechanism by which temperature sensitivity can be encoded.

We proceeded to investigate the molecular basis of thermosensation from a different perspective, by employing SthK, an ion channel from *Spirochaeta thermophila*, as a model protein. This extreme thermophile survives within a temperature range of approximately 40 to 73 °C, with an optimal growth temperature around 65 °C [13], which led us to hypothesize that SthK activity is sensitive to temperature. SthK can be expressed in *E.coli* at high levels, producing large quantities of ion channel sample, lending itself to biophysical and biochemical investigations prohibitive to mammalian ion channels that express only in low amounts [14]. The possibility to isolate the pure ion channel from other cellular components also allows for the manipulation of the environmental lipid composition.

SthK is a bacterial homolog of mammalian cyclic nucleotide-gated (CNG) and hyperpolarization-activated cyclic nucleotide-modulated (HCN) channels, sharing a canonical architecture that includes a K⁺-selective pore, a voltage-sensing domain (VSD), and a cyclic nucleotide-binding domain (CNBD) that responds to cAMP as a partial agonist [14, 15]. SthK displays robust, ligand- and voltage-dependent gating and retains many mechanistic features of its eukaryotic counterparts. Notably, its activity is strongly modulated by lipid composition: anionic phospholipids such as phosphatidic acid (PA) and phosphatidylglycerol (PG) enhance activation, whereas phosphatidylinositol-4,5-bisphosphate (PIP₂) inhibits gating by stabilizing the resting conformation [16, 17]. These properties make SthK a tractable and informative model for dissecting the biophysical basis of thermosensitivity under chemically defined conditions. Motivated by this pronounced lipid sensitivity, we investigated whether the membrane also influences temperature-dependent channel behavior. Although thermosensation has been extensively studied in various ion channels, the role of the lipid environment in shaping thermal responses remains poorly understood. This is particularly relevant given that most thermoreceptors are membrane proteins and thus are embedded in dynamic lipid membranes that may modulate their gating energetics and sensitivity.

Here, we report that the SthK channel is indeed robustly temperature sensitive, with increasing activity at lower temperatures. Unexpectedly, its cold sensitivity is lipid-dependent: amine-containing lipids support activation at low temperatures, whereas anionic lipids abolish it. Given the sharp lipid headgroup-dependence, we investigated the possibility that temperature sensing involves an intersubunit, state-dependent salt bridge—previously shown to be important for gating energetics and sensitive to nearby anionic lipids [16]—as electrostatic interactions such as salt bridges in solution can weaken at low temperatures [18–20]. We indeed found that removal through mutagenesis or weakening of this salt bridge led to elimination or pronounced shifts of temperature sensing, respectively, similar to the effects generated by incorporating different lipids. Lower temperatures, combined with amine-containing lipids, work together to weaken the salt bridge, thereby making SthK open easier with cold temperatures. These findings reveal the molecular mechanism by which a temperature-sensitive channel senses environmental temperature changes.

## Results

### SthK channels are temperature sensitive in specific lipids

To examine the temperature sensitivity of SthK, we purified SthK channels from *E.coli*, reconstituted them in liposomes of different lipid compositions and measured channel activity using a single-mixing stopped-flow assay over a range of temperatures from 10 to 45 °C (Fig. 1a). In this assay, the rate of intraliposomal fluorescence quenching via Tl^+^ entering through open channels is an indicator of SthK activity [21, 22]. Both the protein reconstitution efficiency and the size of proteoliposomes with different lipid compositions were similar across the experimental conditions tested (Extended Data Fig. 1a,b).

**Figure 1.**
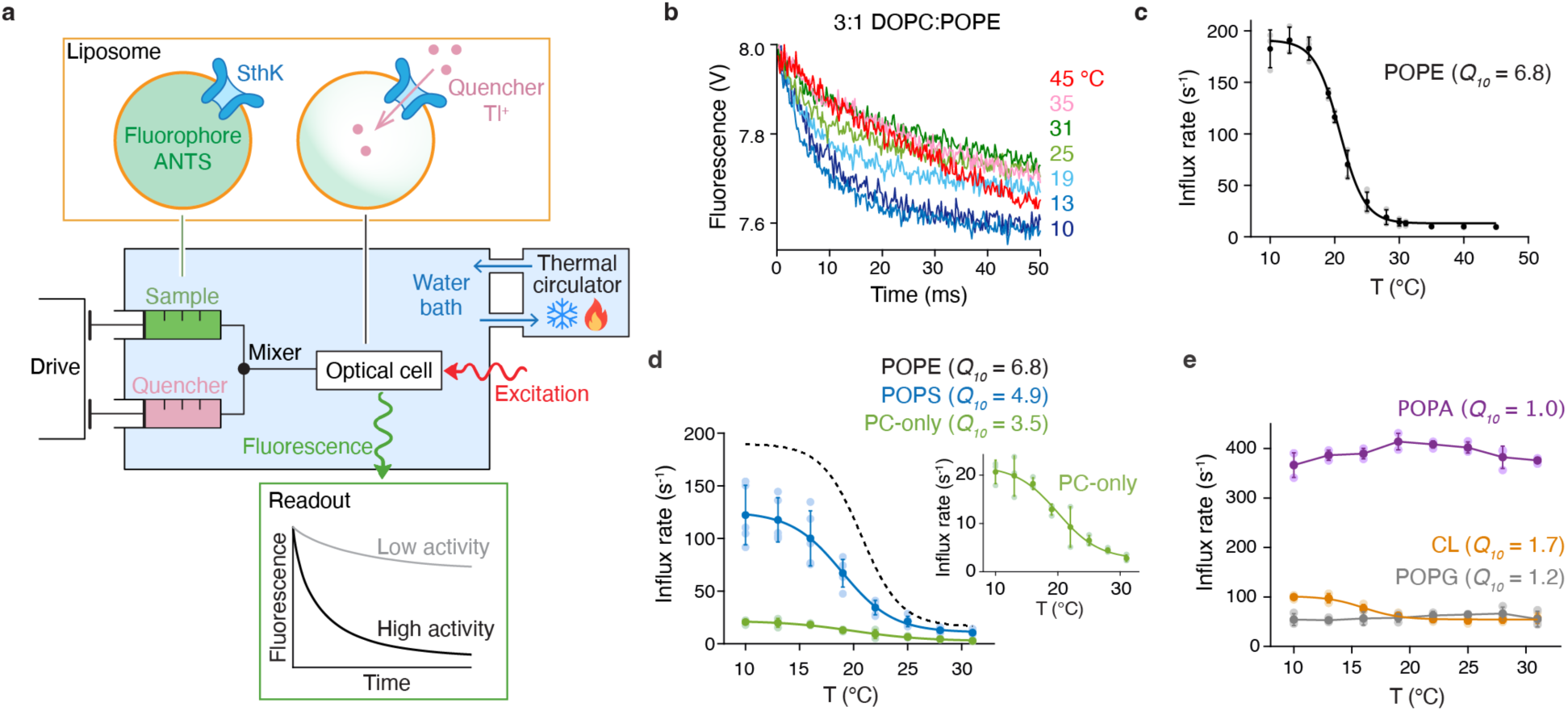
SthK channels are activated by cold temperatures in select lipids. **a**, Scheme of the experimental setup for temperature-controlled channel activity ensemble assays. Syringes with SthK liposomes (green) and Tl⁺ quencher (pink) are equilibrated in a water bath (blue). Upon mixing, Tl⁺ enters via open channels and quenches the ANTS fluorophore (yellow-lined box), reporting activity as kinetics of fluorescence quenching (theoretical activity readout in the green-lined box). **b**, Representative quenching kinetics from stopped-flow assays of SthK reconstituted in 3:1 DOPC:POPE liposomes across 10–45 °C. Only selected traces shown for clarity; full data in Extended Data Fig. 2. **c**, Initial Tl⁺ flux rates from experiments as in **b**. Replicates in light gray and mean ± s.d. in black. Data were pooled from: three independent replicates from 10–31 °C at 3 °C intervals, and three from 20– 45 °C at 5 °C intervals (*n* = 6 at 25 °C, *n* = 3 otherwise). Sigmoidal fit (black line) used the 10–31°C dataset, yielding midpoint temperature (*T_mid_*, °C) = 20.6 ± 0.6 and slope factor (*k*) = 2.1 ± 0.3; the fitted temperature coefficient (*Q_10_*; see Methods and Extended Data Fig. 2) was 6.8 ± 1.4. **d**, **e**, Initial Tl⁺ flux rates for SthK in various lipid environments at different temperatures: PC-only (green) and 3:1 DOPC:X, with X = POPS (blue, *n* = 5), POPA (purple), CL (brown), or POPG (gray); *n* = 3 unless noted. Dashed line in **d** shows PE fit from **c**; inset in **d** highlights PC-only data. Filled circles with error bars are mean ± s.d.; individual data points in lighter shades. Sigmoidal fits yielded *T_mid_* (°C) and slope (*k*), respectively, for: PC-only (20.4 ± 2.2, 2.6 ± 0.5), PS (19.0 ± 1.0, 2.2 ± 0.4), and CL (16.3 ± 1.3, 1.6 ± 0.1); PG and PA were not fitted. Fitted *Q_10_*: PC-only (3.5 ± 0.7); PS (4.9 ± 0.7); PA (1.0 ± 0.1); CL (1.7 ± 0.3); PG (1.2 ± 0.1). All assays were performed with 200 μM cAMP.

The activity of SthK depends on cAMP binding and lipid composition, with higher activity in the presence of anionic lipids [16]. We first tested whether SthK, incorporated in liposomes composed of 3:1 DOPC:POPE lipids, was temperature sensitive over a temperature range from 20 to 45 °C. Surprisingly, instead of increasing with temperature, channel activity rose as the sample was cooled to 20 °C, as indicated by higher influx rates (Fig. 1b,c). We thus extended our investigations to lower temperatures and found that channel activity continued to increase with decreasing temperatures plateauing below 16 °C (Fig. 1b,c). These results indicate that SthK is a cold-sensitive ion channel, with a notable temperature coefficient (*Q_10_*; see Methods for definition) of 6.8 in DOPC:POPE lipids (Fig. 1c, Extended Data Fig. 2).

The above experiments were performed in the presence of cAMP. To investigate whether cooling alone can activate the channel, we tested channel activity in the absence of cAMP after prolonged exposures to low temperatures and found no increase in activity even after extended incubations (Extended Data Fig. 1c). Moreover, in liposomes reconstituted with SthK but without cAMP, we consistently observed negligible activity across all tested temperatures and lipid compositions (Extended Data Fig. 2), suggesting that vesicles did not become leaky at low temperatures. When cAMP was presence, prolonged exposure to low temperatures did not result in activity loss or desensitization either (Extended Data Fig. 1c). To assess whether high temperatures caused irreversible damage to the protein, we re-cooled samples previously heated to 42 °C and remeasured activity (Extended Data Fig. 1d). Channel activity remained unchanged, indicating that the decrease in activity at higher temperatures was not due to protein degradation.

We next asked whether this cold sensitivity depended on the lipid environment. In both DOPC-only liposomes and DOPC liposomes supplemented with POPS (POPC:POPS 3:1), SthK similarly exhibited pronounced cold sensitivity between 13 °C and 28 °C (Fig. 1d), with a midpoint (*T_mid_*) around 20 °C across the tested lipid compositions (See Methods and Extended Data Table 1). Conversely, when DOPC liposomes were supplemented with POPA, POPG, or 18:1 cardiolipin (CL) at a 3:1 ratio, SthK activity displayed little temperature sensitivity over the temperature range investigated (Fig. 1e, Extended Data Fig. 2).

The above results support the conclusion that SthK is temperature sensitive and show that its temperature sensitivity is dependent on the lipid composition. We next investigated the molecular mechanism for thermosensitivity in these channels.

### Lipids with amine-containing headgroups render SthK cold sensitive

We first investigated whether the SthK sensitivity to cAMP changes with temperature by measuring the cAMP *EC₅₀* at 13 °C, 22 °C, and 28 °C. Although a modest increase in *EC₅₀*was observed at 28 °C, the values at 13 °C and 22 °C were statistically indistinguishable (Extended Data Fig. 4), despite a 3-fold difference in channel activity between these temperatures (Fig. 1c). Thus, temperature-dependent changes in SthK sensitivity to cAMP are not a primary driver of the observed thermosensitivity in SthK.

POPE has a melting temperature (*T_m_*) of 25 °C, close to the temperature range where SthK becomes cold sensitive. Although POPE constituted only 25% of the total lipid composition, with the majority being DOPC (*T_m_* = –17 °C), we considered the possibility that lipid phase transition might contribute to temperature sensitivity. To test this, we replaced POPE with DOPE (*T_m_* = – 16 °C, well below the cold-sensitive range) and found that SthK remained cold sensitive, with a *Q_10_* of 8.1 (Fig. 2a), effectively ruling out lipid phase transition as the underlying cause.

**Figure 2.**
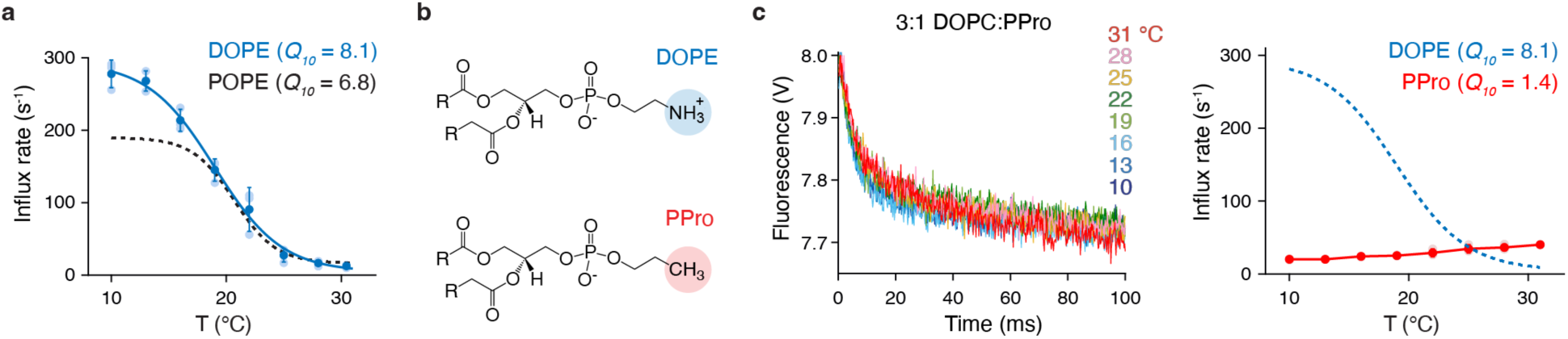
Amine-containing lipid headgroup is essential for cold sensing in SthK. **a**, Initial Tl^+^ flux rates for SthK in 3:1 DOPC:DOPE liposomes (blue, *n* = 3) compared with 3:1 DOPC:POPE (reproduced as a dashed line from Figure 1c). Filled symbols are mean ± s.d.; individual data points in lighter shades. Sigmoidal fit (solid line) yielded *T_mid_* = 19.3 ± 0.9 °C and slope factor *k* = 2.4 ± 0.3. The corresponding temperature coefficient (*Q_10_*, see Extended Data Fig. 2) was 8.1 ± 1.0. **b**, Structures of DOPE and 18:1 PPro, with identical tails omitted for clarity. The only difference—the presence of a primary amine (NH₃⁺) in DOPE versus a methyl group (CH₃) in PPro—is highlighted. **c**, Representative quenching kinetics (left) and initial Tl⁺ flux rates (right, red symbols, *n* = 3) of SthK in 3:1 DOPC:PPro at different temperatures. Filled symbols are mean ± s.d., with individual experiments shown as lighter-shaded circles. The fitted *Q_10_* for PPro was 1.4 ± 0.1. Dashed line shows 3:1 DOPC:DOPE flux rates from **a** for reference. All assays were performed with 200 μM cAMP.

We next asked whether specific chemical features of the lipid headgroups contribute to SthK cold sensitivity. PC, PS and PE all have amine-containing headgroups (Extended Data Fig. 5). To investigate whether the amine group is essential for SthK cold sensitivity, we reconstituted SthK in DOPC supplemented with 18:1 phosphatidylpropanol (PPro, Fig. 2b). DOPE and PPro have similar structures that differ by only one atom; PPro has a methyl group in place of the amine in PE. In PPro-containing liposomes, the cold sensitivity of SthK is completely abolished (Fig. 2c), indicating that the amine group is indeed essential for the cold sensitivity of SthK.

### Structure of SthK in PE-containing lipid nanodiscs

To explore the interaction between the lipid amine and SthK at lower temperatures, we reconstituted SthK in the presence of cAMP in lipid nanodiscs containing 1:1 DOPC:DOPE lipids and plunge-froze the grids at 6 °C for single-particle cryo-EM analysis. Two distinct conformations were identified (Extended Data Fig. 6 and Extended Data Table 2), both very similar to previously reported closed SthK structures [15, 16]. These two structures are highly similar to each other, with a Cα RMSD of 0.41 Å for the transmembrane domain (TMD) and 0.42 Å for the cyclic nucleotide-binding domain (CNBD) when aligned separately, and both exhibit a closed pore consistent with a non-conducting state (Fig. 3a-b), despite reconstitution into nanodiscs containing PE and grid preparation at low temperature intended to open the channel. This was likely due to the intrinsically low open probability (*P_o_*) of SthK under these conditions, consistent with previous single-channel recordings performed at room temperature, showing <0.1% *P_o_* at 0 mV in 25% PE [16], as well as our own stopped-flow results indicating low activity in PE at room temperature. Although lowering temperature under these conditions increases activity by several fold, the overall open population remains still too low (∼1%) for structural identification.

**Figure 3.**
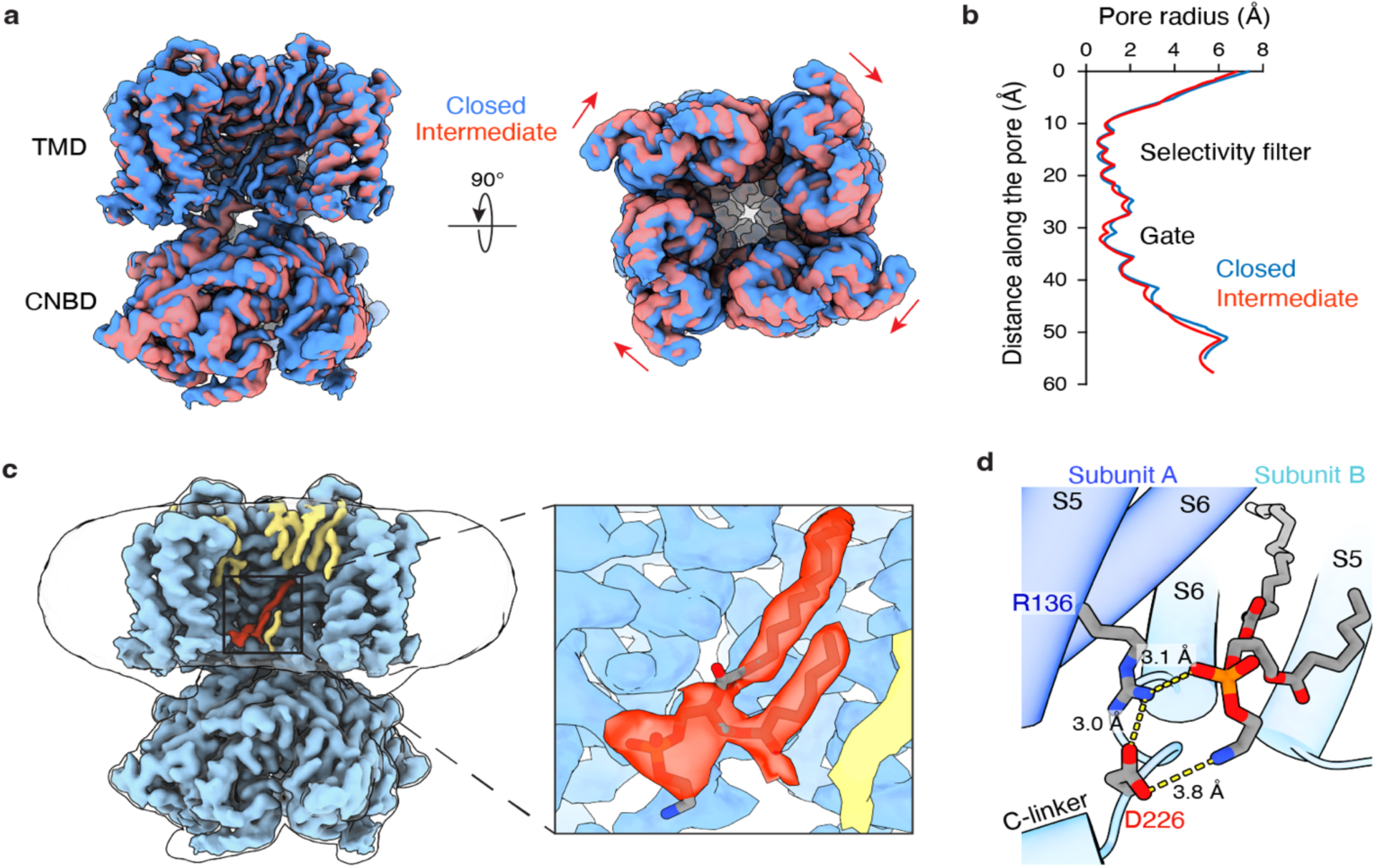
Structures of SthK in complex with PE lipids. **a**, Cryo-EM maps of the closed (blue) and intermediate (pink) states, aligned by their transmembrane domains (TMDs). **b**, Pore radius profiles calculated using HOLE. **c**, Density map of the closed state with protein in blue, most lipid densities in yellow, and a well-defined lipid density with both head group and two tails discernible in the inner leaflet highlighted in red. The inset is a zoomed-in view of the red density, modeled as a PE lipid. **d**, Atomic model showing a PE molecule headgroup within interaction distance of the intersubunit salt bridge between R136 on the S5 helix (subunit A, dark blue) and D226 on the S6 helix (subunit B, light blue). Yellow dashed lines indicate potential salt bridge formations.

The difference between the two structures lies in a minor relative rotation between the CNBD and the TMD (Fig. 3a), which led a small changes in the distance that separates Arg136 and Asp226 (Extended Data Fig. 7a), two residues that are part of an intersubunit salt bridge (R136–D226) previously reported to be involved in stabilizing channel closed states. This salt bridge is state-dependent: the R136–D226 distance is short in the closed conformation and large during the upward-rotation movement of the CNBD, a key step toward channel opening [16]. The SthK conformation with the shorter salt bridge distance and stronger interaction will be referred to as the closed state. Since the larger R136–D226 distance reflects a loosened intersubunit constraint, primed for pore opening, we call this state an intermediate state, to reflect its conformational position along the gating trajectory.

Both SthK conformations display lipid density near this salt bridge (Fig. 3c and Extended Data Fig. 7b). In the closed conformation, the lipid density is distinct with two carbon tails and a continuous lipid head group, and we modeled it as a PE molecule (Fig. 3c,d). Its amine group (positively charged) is within interaction distance of the side chain of D226 (negatively charged), in a position to electrostatically weaken the salt bridge. This is reminiscent of the previously reported interaction between the negatively charged phosphate of a PA lipid binding at the same location and the positively-charged R136, which was shown to effectively eliminate the R136– D226 salt bridge interaction [16]. Since salt bridges have been previously reported to vary in strength with temperature, with cold temperatures reportedly weakening them––a phenomenon primarily linked to reduced desolvation penalty at lower temperatures [18–20, 23]––we next investigated whether this intersubunit salt bridge serves as the temperature sensor of SthK, and whether its weakening by the lipid amine headgroup is the main cause of SthK’s cold sensitivity in the presence of amine-containing lipids.

### Intersubunit salt bridge is the cold-sensor of SthK

If the intersubunit salt bridge serves as the cold sensor of SthK, its removal, by mutating Asp226 to Asn (D226N) should eliminate temperature sensitivity. We indeed found that cold sensitivity was completely abolished in SthK D226N in PE-containing liposomes, and the mutant channel activity was high, similar to that observed in SthK WT in PE-containing liposomes at low temperatures (Fig. 4a). However, we cannot exclude the possibility that SthK D226N is constitutively active due to the loss of the salt bridge, thereby masking any temperature-elicited activity increases. Hence, we next sought a milder disruption of the salt bridge as opposed to breaking it completely.

**Figure 4.**
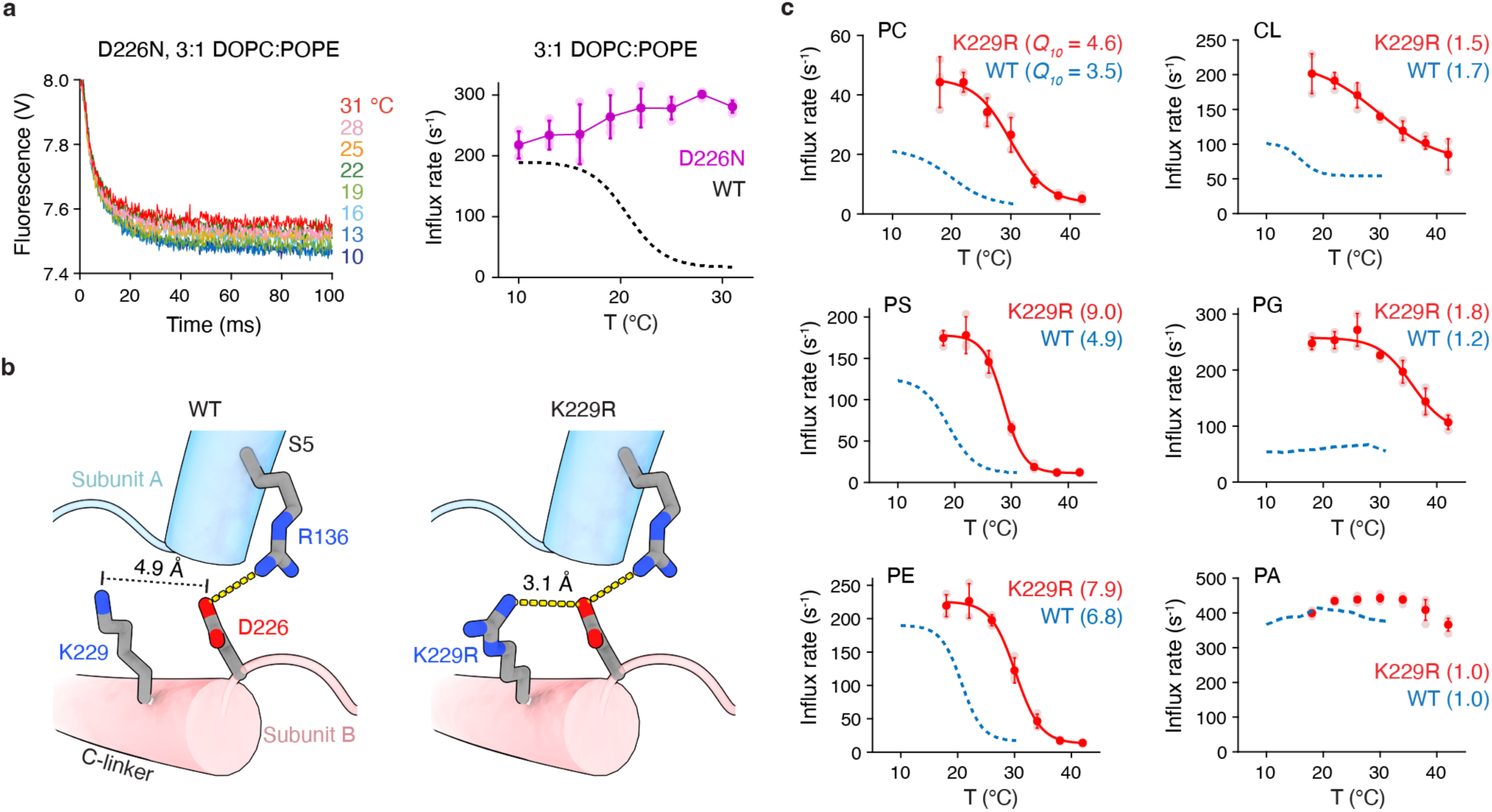
The intersubunit salt bridge is the cold sensor of SthK. **a**, Quenching kinetics (left) and initial Tl^+^ flux rates (right) from stopped-flow flux assays for SthK D226N, reconstituted in 3:1 DOPC:POPE. Dashed line shows WT data (from Fig. 1c) in the same conditions, for comparison. Dark purple symbols with error bars represent mean ± s.d. from 3 biological replicates. Light shaded symbols are the individual data points. **b**, Detail from the structure of WT SthK (PDB: 7tj5) on the left of the intersubunit salt bridge between R136 and D226, highlighting the neighboring residue K229, located one helical turn below D226. The K229R SthK model (right), generated by Mutagenesis Wizard in PyMOL, displays the lowest-strain rotamer for the R229 side chain. Yellow dashed lines indicate distances compatible with potential salt bridge formation. **c**, Initial Tl^+^ flux rates for SthK K229R (red symbols with error bars represent mean ± s.d.) reconstituted in different lipid compositions over a range of temperatures. From top to bottom and left to right: PC-only (*n* = 3); 3:1 DOPC:X, where X = POPS (*n* = 3), POPE (*n* = 3), CL (*n* = 3), POPG (*n* = 3), or POPA (*n* = 4). Data showing temperature-associated changes were fitted with sigmoidal curves (solid red lines). Fitted midpoint temperatures (*T_mid_*, °C) and slope factors (*k*) for SthK K229R, respectively: PC-only (29.4 ± 2.3, 2.7 ± 0.9), PS (28.5 ± 0.5, 1.9 ± 0.2), PE (30.1 ± 0.6, 2.1 ± 0.4), CL (29.3 ± 1.1, 4.1 ± 0.6), PG (33.5 ± 0.6, 3.8 ± 1.7); PA: not fitted due to lack of a clear sigmoidal transition. Fitted temperature coefficients (*Q_10_*, see Extended Data Fig. 2) for SthK K229R: PC-only (4.6 ± 0.9), PS (9.0 ± 1.0), PE (7.9 ± 0.6), CL (1.5 ± 0.2), PG (1.8 ± 0.1), PA (1.0 ± 0.1). WT data in corresponding lipid compositions are shown as blue dashed lines (from Fig. 1c–e), for comparison. All assays were performed with 200 μM cAMP.

Lys229 (K229) is one helical turn away from D226 (Fig. 4b). In SthK WT, the side chain of K229 is relatively far from the side chain of D226 (4.9 Å). We hypothesized that a longer arginine side chain at the same position (K229R) may get within interaction distance of D226, and this way weaken the R136–D226 salt bridge. Such manipulation of the temperature sensor is predicted to perturb rather than eliminate the temperature sensitivity of SthK.

We examined the activity of SthK K229R in liposomes containing only PC or supplemented with PS or PE (Fig. 4c, left). The activity of SthK K229R was overall higher than WT, and the midpoint temperature (*T_mid_*) in each of these lipid compositions was shifted by approximately 10 °C towards an increased sensitivity to cold temperatures (Fig. 4c, Extended Data Table 1). This is consistent with our hypothesis that the salt bridge in the K229R mutant is weaker, which leads to a destabilization of the closed state, enabling channel opening at less cool temperatures than WT. Interestingly, in the presence of anionic lipids such as PG or CL, where SthK WT showed no temperature sensitivity, SthK K229R displays a large increase in activity at cold temperature (Fig. 4c, right). We attribute this to a weakening of the intersubunit salt bridge, jointly caused by the arginine substitution (K229R) and the presence of anionic lipids [16]. Together, these factors destabilize the closed state and enhance channel activity at lower temperatures. In the presence of PA, no observable change in activity was noted, likely because the disruption of the intersubunit salt bridge by the small head group of PA is already maximal even in SthK WT [16]. Taken together, our results showed that the intersubunit salt bridge R136– D226 serves as the cold sensor of SthK, and perturbation of this salt bridge with either mutations or via the type of lipid bound near the salt bridge alters the temperature sensitivity of SthK.

## Discussion

In this study, we report the mechanism by which the SthK ion channel senses temperature changes (Fig. 5a). We found that SthK is a cold-sensing ion channel, exhibiting higher activity at lower temperatures. The cold sensitivity of SthK is lipid-dependent, showing particular sensitivity in the presence of amine-containing lipids such as PC, PS, or PE. Through structural and functional studies, we have demonstrated that the intersubunit salt bridge R136–D226 acts as the temperature sensor of SthK. Our results show that the intersubunit salt bridge is weakened by lipid amines and further compromised by low temperatures, resulting in channel opening at temperatures below 30 °C. Among amine-containing lipids, SthK shows particularly strong cold activation in the presence of PE compared to PC or PS. This may reflect the smaller and less bulky geometry of the PE head group (Extended Data Fig. 5), which could fit more snugly in the binding site and thus more effectively destabilize the salt bridge and enhance cold sensitivity.

**Figure 5.**
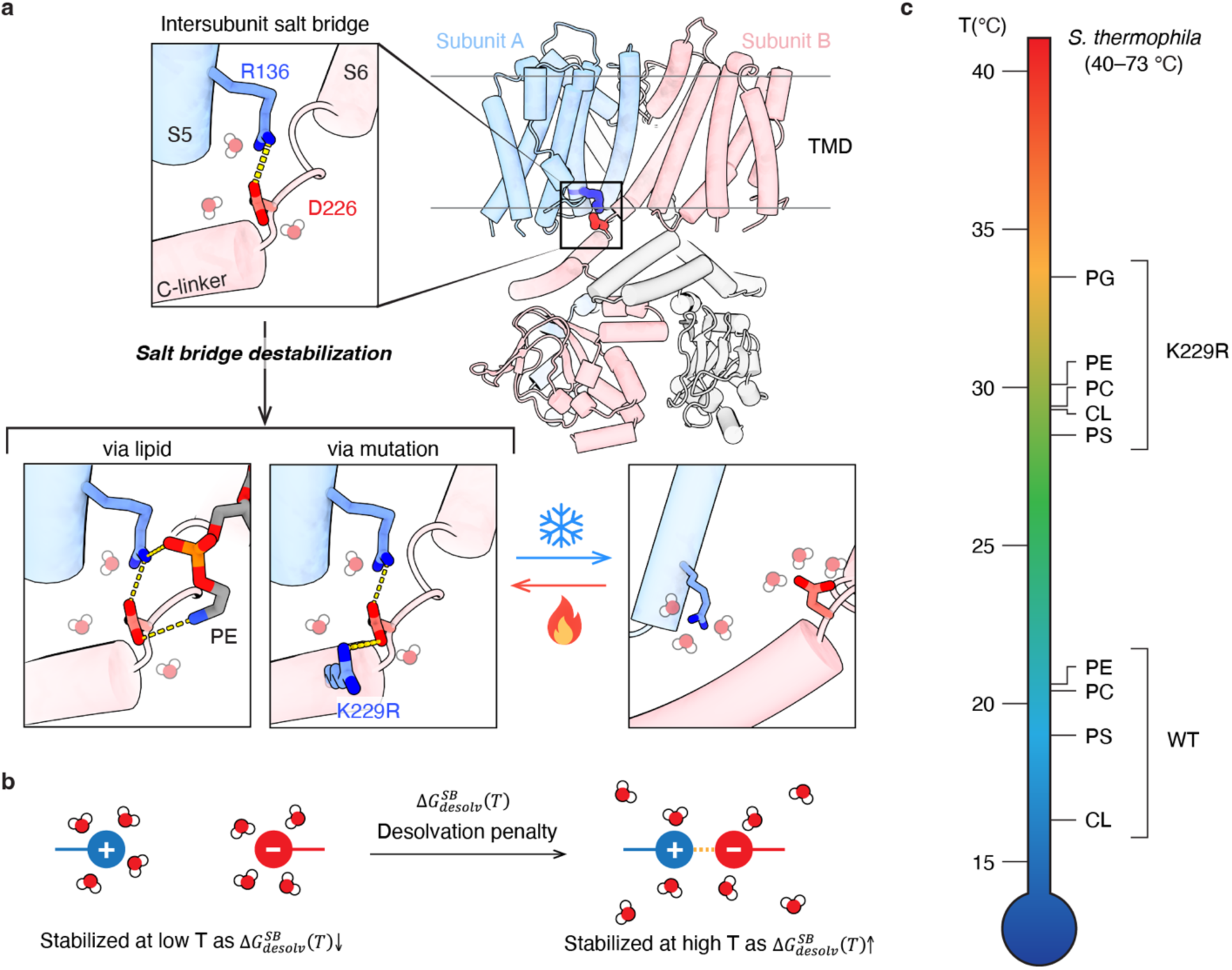
Model for temperature-dependent gating via modulation of salt bridge strength. **a**, An intersubunit salt bridge between R136 and D226 stabilizes the channel in closed states. Disruption of the strength of the salt bridge by either surrounding lipid headgroups (e.g., PE) or mutations such as K229R weakens the bridge sufficiently so that it becomes sensitive to temperature changes in the range tested here. The model of the broken salt bridge on the right is from the previously published open structure of SthK (PDB: 7tj6). Charge-bearing atoms of R136, D226, and K229R are colored dark blue (positive) and red (negative), respectively. Yellow dashed lines indicate potential salt bridge formation. **b**, Conceptual illustration showing how the temperature dependence of desolvation penalty upon salt bridge (SB) formation, 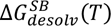, governs salt bridge stability. At lower temperatures, unpaired charged residues are stabilized by stronger hydration, resulting in a smaller 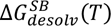, which disfavors salt bridge formation. At higher temperatures, hydration weakens, increasing 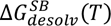 and promoting the formation of the salt bridge. **c**, Cartoon highlighting the wide temperature range of midpoint temperatures (*T_mid_*) achievable for cold activation in SthK WT and K229R in different lipid environments. This highlights the ease of tuning temperature sensitivity of channels via this mechanism by either lipids with different headgroups bound near the salt bridge or by having select residues located within interaction distance of the salt bridge.

The mechanism proposed here may extend to other cold thermoreceptors that feature state-dependent salt bridges. In TRPM8, a well-characterized cold thermoreceptor, an intersubunit salt bridge between Arg851 and Asp866 in mouse TRPM8 has been shown to form in the closed state and break upon agonist-induced channel opening [24]. Additionally, this salt bridge is involved in PIP₂ binding. The convergence of state dependence and lipid sensitivity at this site raises the possibility that TRPM8 may share a similar lipid-modulated cold-sensing mechanism with SthK. Although cold and heat thermoreceptors exhibit opposite trends in response to temperature changes—cold-sensing thermoreceptors become more active as temperature decreases, while heat-sensing thermoreceptors become less active—the underlying principles governing salt bridge dynamics may be shared. For instance, heat-activation in thermoreceptors might involve the strengthening of a strategically positioned salt bridge, stabilizing the open state and promoting channel activity at elevated temperatures.

How can weakening of a salt bridge function as a cold sensor for an ion channel, ultimately leading to channel opening at lower temperatures? The R136–D226 salt bridge is located at the water–lipid interface, rendering it sensitive not only to the chemical nature of the lipid headgroups but also to the surrounding solvent environment (Extended Data Fig. 7c). Prior studies have suggested that temperature-dependent changes in solvent dielectric constant may weaken salt bridges at lower temperatures, as electrostatic attraction decreases with increasing dielectric constant [23]. However, such effects are modest within the temperature range of our system (e.g., 76.5 at 30 °C to 83.8 at 10 °C for water [25]) and likely contribute only marginally. In contrast, the desolvation penalty—the energetic cost of dehydrating charged side chains to form a salt bridge— is more strongly temperature-dependent (Fig. 5b) [18–20]. Previous studies have shown that hydration is stronger at lower temperatures, leading to a higher desolvation penalty and greater energetic cost for solvent removal, which in turn disfavors salt bridge formation [18–20].

Together, these solvent and lipid effects may act in concert to destabilize the salt bridge, priming the channel for opening under cold conditions. Notably, the state-dependent Arg851– Asp866 salt bridge in mouse TRPM8 is also situated near the membrane interface, raising the possibility that it too may be subject to modulation by lipid and solvent effects [24].

While the process of salt-bridge weakening at lower temperatures has been previously reported, most of these reports are based on in silico studies [18–20, 26–28]. Experimental evidence is limited, and to the best of our knowledge, only one other study, on low temperature favoring tetramer to dimer dissociation of a L-xylulose reductase (XR) due to the weakening of the intersubunit salt bridge, has demonstrated this phenomenon [29]. The tendency of salt bridges to weaken at lower temperatures is also reflected in nature. Proteins from thermophilic organisms typically contain more salt bridges than those from mesophilic species, which enhances thermal stability by reinforcing intramolecular interactions at high temperatures [30–34]. However, this same feature often makes thermophilic proteins less stable at ambient temperatures, as the higher number of salt bridges, while stabilizing at elevated temperatures, can become marginal, or even destabilizing, when electrostatic interactions weaken at lower temperatures [33, 34].

The intersubunit salt bridge between R136 and D226 was previously shown to also be susceptible to disruption by anionic lipids [16]. PA robustly activates SthK across a broad temperature range, rendering the channel highly active, which could mask any further temperature-induced increases. PG and CL also show no temperature dependence but they increase channel activity to a much lesser extent than PA [16], which would argue against high channel activity masking the temperature effect. In addition, these lipids were proposed to destabilize the salt bridge via their charged headgroup, similar to the amine-containing lipids, so why don’t they also confer temperature sensitivity to SthK? A previous computational study reported that PG-containing membranes not only promote stronger arginine–lipid headgroup interactions than PC-only bilayers but also exhibit a localized enrichment of cationic counter-ions at the membrane interface, enhancing local electrolyte screening [35]. We speculate that anionic lipids such as PG or CL give rise to a more electrostatically buffered interfacial environment, where desolvation penalty associated with salt bridge formation are attenuated. In such environments, hydration shells around charged residues may already be partially disrupted by elevated local ion concentrations, thereby reducing the entropic gain upon salt bridge formation and diminishing its temperature sensitivity.

The salt bridge–weakening mutation K229R results in a higher overall *Q_10_* in the presence of amine-containing lipids (PC, PS, or PE), and markedly increased channel activity at lower temperatures in the presence of anionic lipids such as PG and CL although the *Q_10_*values remain relatively low (*Q_10_*= 1.8 and 1.5, respectively). Notably, SthK exhibits an inverse thermal response—higher activity at lower temperatures—which contrasts with the temperature dependence of ion diffusion, where mobility increases with temperature (*Q_10_* = 1.3–1.5 for diffusion of ions alone) [36]. As a result, the observed *Q_10_* for SthK may underestimate the intrinsic temperature sensitivity of its gating, since the reduced rate of ion diffusion at lower temperatures partially offsets the increase in channel activity under those conditions.

The temperature sensor identified here for SthK—a state-dependent intersubunit salt bridge—is highly conserved in HCN channels (Extended Data Fig. 8) at both the sequence and structural levels, suggesting that they may employ a similar lipid-dependent cold-sensing mechanism. While HCN channels are not considered primary cold thermoreceptors, accumulating evidence points to their modulatory role in cold-sensitive neurons: for instance, inhibition or genetic deletion of HCN channels reduces behavioral responses to cooling in mice [37, 38], and oxaliplatin-induced cold hypersensitivity can be alleviated by the HCN inhibitor ivabradine [39]. Whole-cell recordings of human HCN2 expressed in HEK293 cells showed reduced activity at lower temperatures [40], which appears inconsistent with cold-sensing properties. However, our findings with SthK suggest that thermosensitivity is highly contingent on the surrounding lipid environment and the complex and heterogeneous membrane composition of HEK293 cells may mask a thermosensitive behavior of HCN channels. This raises the possibility that, in select lipid environments HCN channels may indeed respond to temperature. Future studies may offer broader insights into how membrane composition shapes thermal responsiveness in ion channels and open avenues for targeting temperature-sensitive channels in physiological and pathological contexts.

The larger increase in activity at lower temperatures and the considerably higher temperature midpoint displayed in SthK K229R compared to the extent to which lipid composition modulates WT SthK may reflect the inherent flexibility of bound lipids and their limited ability to destabilize the salt bridge as much as the consistently positioned K229R side chain. This way, the presence of the K229R side chain with its more persistent destabilizing effect on the salt bridge, promotes channel activation at less cool temperatures. While these observations support a model in which salt bridge destabilization contributes to temperature sensitivity, the mechanism proposed here may be oversimplified. Mutation and lipid likely modulate salt bridge stability through distinct physical principles, including differences in spatial persistence and interaction dynamics, and their potential impact on desolvation penalty upon salt bridge formation, and further studies will be required to disentangle these potentially divergent effects.

The temperature-sensitive range of wild-type SthK (10–30 °C) falls below the survival temperature range of *Spirochaeta thermophila* (40–73 °C) [13], the bacterium from which SthK was identified. Since the lipid composition of this organism remains largely uncharacterized, it is possible that in its native lipid or cellular environment, the state-dependent salt bridge of SthK responds within a temperature range closer to its survival range. This can be parallelled to the substantial *T_mid_* shift observed in the K229R relative to wild-type SthK in most lipid compositions tested (Fig. 5c). Notably, K229R exhibits a *T_mid_* of 33.5 °C in PG-containing liposomes, approaching the survival range of *S. thermophila*. Altogether, these findings suggest that SthK’s thermal sensitivity is shaped not only by the residues found in the vicinity of the salt bridge but also by its surrounding chemical and physical environment, which may differ substantially under physiological conditions.

The ability of SthK and, presumably, other temperature-sensitive ion channels to respond to temperature may therefore arise from a finely tuned balance between structural elements and the membrane environment. Not only must the salt bridge possess an intermediate strength— neither too strong to resist thermal modulation nor too weak to contribute meaningfully to gating— but its sensitivity can also be easily masked by the surrounding lipid composition. For example, lipid environments lacking destabilizing headgroups, such as the amine-containing lipids examined here, may stabilize the salt bridge and obscure any thermally induced gating effects. Conversely, excessive destabilization could prevent the salt bridge from forming altogether, eliminating its contribution to channel function. Thus, as with the Goldilocks principle, only through the combined interplay of salt bridge properties and membrane context can the system reach the “just right” conditions needed for temperature sensitivity.

### Conclusion

Here, we reveal that SthK ion channel senses temperature changes through a lipid-modulated intersubunit salt bridge (R136–D226) located at the membrane interface. We demonstrate that weakening of this salt bridge at lower temperatures, particularly in the presence of amine-containing lipids, promotes channel opening and cold activation. The temperature-sensing gating via salt bridge interaction strength modulation may only emerge under finely tuned structural and environmental conditions—an energetic “Goldilocks zone” where the interaction is neither too strong nor too weak. Identifying such a mechanism in SthK not only uncovers a novel form of lipid-dependent thermosensitivity but also provides a framework for exploring similarly constrained temperature-sensing strategies in other ion channels.

## Methods

### Expression and purification of SthK

Protein expression and purification was performed as previously described [14]. *E. coli* cells were grown in LB media at 37 °C, supplemented with ampicillin. Upon reaching OD_600_ of 0.4, the temperature was lowered to 20 °C for further growth until OD_600_ reached 0.8. Protein expression was initiated by adding 0.5 mM isopropyl-β-d-thiogalactoside (IPTG). Cells were harvested after 12–14 h through centrifugation at 6000g for 10 min at 4 °C. The cell pellet was resuspended in breaking buffer (50 mM Tris, 100 mM KCl, pH 8) with 1 mg of lysozyme (Sigma-Aldrich), 1 mg of DNase I (Sigma-Aldrich), 85 μg ml^-1^ PMSF (Roche), Leupeptin (0.95 μg ml^-1^, Roche), Pepstatin (1.4 μg ml^-1^, Roche), and ultra cOmplete protease inhibitor (Roche), and further disrupted by sonication. To extract SthK from cell membrane, lysate was supplemented with 30 mM DDM (n-dodecyl-β-d-maltopyranoside, Anatrace) and incubated for 1.5 h at 4 °C with constant agitation. The insoluble fraction was removed by centrifugation at 37,500g for 50 min at 4 °C. The supernatant was filtered through a 0.22 μm membrane and loaded onto a 5 ml Hitrap column (Cytiva) charged with Co^2+^. The column was washed with 75 ml of wash buffer (20 mM HEPES, 100 mM KCl, 100 µM cAMP, 30 mM imidazole, 0.5 mM DDM, pH 7.8) and SthK was eluted with 10 ml of elution buffer (20 mM HEPES, 100 mM KCl, 100 µM cAMP, 300 mM imidazole, 0.5 mM DDM, pH 7.8). The protein was concentrated to about 15 mg ml^-1^ and further purified by gel filtration on a Superdex 200 Increase 10/300 GL column (Cytiva) in 20 mM HEPES, 100 mM KCl, 0.5 mM DDM, pH 7.4. The SthK fraction was pooled, concentrated to about 10 mg ml^-1^, flash frozen in liquid nitrogen, and stored at −80 °C for future use. All purification steps were performed at 4 °C.

Mutations in SthK were introduced by site-directed mutagenesis using Q5 polymerase (New England Biolabs) and the following primers: D226N-forward: CTC GTC TCG AAA CTG AAT GCC GCA AAG CTC CTC, D226N-reverse:GAG GAG CTT TGC GGC ATT CAG TTT CGA GAC GAG, K229R-forward: AAA CTG GAC GCC GCA AGG CTC CTC CAC, and K229R-reverse: CTC CCT GTG GAG GAG CCT TGC GGC GTC. Due to lower expression levels observed in SthK mutants, *E. coli* C43 cyaA^-^ cells were used for protein expression [41, 42]. Cells were grown in terrific broth media at 37 °C until reaching an OD_600_ of 1, followed by induction with 0.5 mM IPTG at 20 °C for 12–14 h. The subsequent protein purification steps were carried out as for SthK WT.

### Preparation of proteoliposomes for stopped-flow assay

For single-mixing stopped-flow Tl^+^ influx assay, purified SthK was reconstituted into large unilamellar vesicles encapsulated the fluorophore ANTS (8-aminonaphthalene-1,3,6-trisulfonic, Life Technologies) as previously described [21, 22, 43]. Initially, 10 mg of lipids in chloroform were dried to a thin film in a glass tube under a gentle N_2_ gas stream, followed by overnight drying under vacuum. The dried lipids were rehydrated in 743 μl reconstitution buffer (10 mM HEPES, 140 mM KNO_3_, pH 7.4) supplemented with 50 mM CHAPS detergent (3-((3-cholamidopropyl) dimethylammonio)-1-propanesulfonate). The resulting solution was sonicated until transparent. Subsequently, 300 μg of purified SthK and 371 μl of ANTS solution (reconstitution buffer supplemented with 75 mM ANTS, pH 7.4) were added and incubated for 10 minutes at room temperature. An additional 667 μl ANTS solution was added to reach a final concentration of 25 mM. Detergent was removed by addition of 667 mg of BioBeads (SM-2, BioRad) as a 50% slurry in reconstitution buffer, followed by a 3-hour incubation under constant agitation at 21 °C. The supernatant was transferred to a new glass tube and stored at 13 °C overnight. The next day, liposomes were briefly sonicated in a water bath sonicator, extruded through a 100 nm filter (mini extruder, Avanti Polar Lipids), and external ANTS was removed passing through a PD-10 desalting column (Cytiva) pre-equilibrated with reconstitution buffer. To eliminate any trace amount of external ANTS, only the initial 3 ml of the eluted fraction from desalting column was collected. The eluted fraction was further diluted with reconstitution buffer to a final lipid concentration of 0.227 mg ml^-1^ for the following single-mixing stopped-flow experiments.

### Single-mixing stopped-flow Tl^+^ influx assay and data analysis

Stopped-flow Tl^+^ influx experiments were performed in a single-mixing fashion with fluorescence detection (Applied Photophysics). Before recording, SthK was activated by mixing the proteoliposomes with cAMP to reach a final concentration of 200 μM. The resulting activated proteoliposomes were then mixed with quenching buffer (10 mM HEPES, 90 mM KNO_3_, 50 mM TlNO_3_, 200 µM cAMP, pH 7.4) in the stopped-flow device, and the fluorescence signal was recorded. Fluorescence measurements were performed with an excitation wavelength of 360 nm, utilizing a 420 nm high-pass filter, capturing 5,000 data points over a 1-second interval. At least ten repeats were performed for each temperature during a single experiment. Kinetics were inspected visually and mixing artifacts were sorted out. Fluorescence quenching data were analyzed as previously described [21]. Each quenching kinetics was analyzed using a stretched exponential:

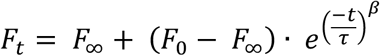

where *F_t_*, *F_∞_*, *F_0_* represent the fluorescence a time *t*, the final fluorescence, and the initial fluorescence, respectively. The variables *t*, ***τ***, and *β* denote time (in s), the time constant (in s), and the stretched exponential factor, respectively. The utilization of stretched exponential accounts for the size distribution of liposomes and the presence of varying numbers of active channels per liposome. The Tl^+^ influx (*k_t_*) at 2 ms for each temperature was calculated using equation:

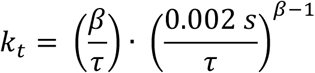

Subsequently, the rates for each temperature were averaged, and the standard deviations were determined. Every experiment was replicated a minimum of three times, and the acquired rates were then averaged, accompanied by corresponding standard deviations. The precise number of repetitions (*n*) is specified in the figure legends, and each repetition covered the entire temperature range (indicated in figure legends), starting from the lowest temperature to the highest temperature. Data acquisition was executed through ProDataSX (Applied Photophysics), and subsequent analysis was conducted in MATLAB (MathWorks).

To determine the temperature sensitivity of SthK in different lipid compositions, the temperature coefficient (*Q_10_*) was calculated by linear fitting of the log-transformed, normalized Tl^+^ influx rates over the range of steepest temperature dependence, as visually identified from the log-transformed influx plot. The absolute value of the slope (|*m*|) was then used to compute *Q_10_* as:

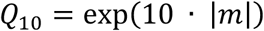

This definition ensures all *Q_10_* values are ≥1 to facilitate intuitive comparison.

To assess the temperature-sensitive range of SthK in different lipid compositions, the midpoint temperature (*T_mid_*) is determined by using a sigmoidal curve fitting:

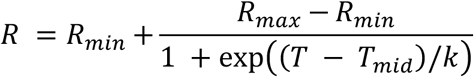

where *R_min_* and *R_max_* represent the minimal and maximal influx rates, respectively, and *k* is the slope factor. The midpoint temperature *T_mid_* provides a central reference point within the temperature-sensitive range, indicating where the influx rate changes most rapidly.

### Nanodisc reconstitution for cryo-EM structural determination

To eliminate endogenous lipids that may tightly bind to SthK, the buffers used for immobilized metal ion affinity chromatography and size-exclusion chromatography were supplemented with 0.02 mg ml^-1^ DOPE, facilitating efficient lipid exchange. The membrane scaffold protein MSP1E3 was prepared according to established methods [44] and concentrated to 15 mg ml^-1^ in reconstitution buffer (10 mM HEPES, 100 mM KCl, pH 7.4). A lipid stock solution of DOPC:DOPE at a 1:1 ratio was prepared at 100 mM, solubilized in reconstitution buffer supplemented with 200 mM sodium cholate (Anatrace). The nanodisc reconstitution was conducted in a 250 μl volume, aiming for a final SthK concentration of 100 μM. SthK, MSP1E3, and lipids were mixed at a molar ratio of 1:1.1:70 in reconstitution buffer supplemented with 3 mM cAMP, followed by a 20-minute incubation at room temperature. Detergent removal commenced with the addition of 100 mg BioBeads (BioRad) equilibrated in reconstitution buffer. Samples were gently agitated and incubated for 2 hours at 4 °C, with subsequent transfer of the supernatant to a new tube containing 100 mg BioBeads. Further incubation at 4 °C overnight under gentle agitation was performed. The pooled supernatant was combined with a wash of the BioBeads using a total of 250 μl reconstitution buffer. The 500 μl sample was filtered through a Spin-X column (Costar) and loaded onto a Superose 6 Increase 10/300 GL gel filtration column (Cytiva) equilibrated in reconstitution buffer supplemented with 200 μM cAMP. The peak corresponding to nanodiscs containing SthK was collected and concentrated to approximately 8 mg ml^-1^ (1 absorbance (A280) = 1 mg). The final sample was supplemented with cAMP to a final concentration of 3 mM and fluorinated Fos-choline-8 (Anatrace) to a final concentration of 3 mM in reconstitution buffer.

### Cryo-EM grid preparation and data collection

UltrAu-Foil R1.2/1.3 300-mesh gold grids (Quantifoil) were glow-discharged for 80 s. A volume of 3.5 μl of ice-cold sample was applied to the grid, followed by a 30 s incubation at 6 °C and 100% humidity. Subsequently, the grid was blotted for 2.5 s with 0 blot force and plunge-frozen in liquid ethane using a Vitrobot Mark IV (FEI, Thermo Fisher).

Data acquisition was carried out using a Titan Krios microscope (FEI, Thermo Fisher) operated at 300 kV, equipped with a GatanK3 imaging system at ×105,000 nominal magnification. The calibrated pixel size was set at 0.4125 Å (super resolution), and a 20 eV energy filter slit width was used throughout the collection. Videos were recorded at a dose rate of 26.89 e^−^/Å^2^ per s, with a total exposure of 1.80 s, accumulating a dose of 48.40 e^−^/Å^2^ using Leginon [45]. Frames were recorded every 0.05 s for a total of 40 frames per micrograph at a nominal defocus range of 0.4 to 2.5 µm.

### Cryo-EM data processing

Cryo-EM data processing was carried out using Relion-4.0 [46]. A total of 7,118 movies were imported, motion-corrected, and binned three times. CTF estimation without dose-weighting of the micrographs was performed with CTFFIND 4.1 [47]. Subsequently, 1,383,889 particles were picked using crYOLO [48] after fine-tuning the general model for SthK and extracting them with a 256-pixel box size. Junk particles were removed through 2D classification, followed by 3D refinement using an ab-initio SthK model as a reference in C1 before further sorting particles by 3D classification without alignment. The selected particles underwent multiple rounds of CTF refinement, Bayesian polishing, and 3D refinement in C4, as no asymmetry was observed. The closed and intermediate states were then selected after an additional round of 3D classification in C4 without alignment.

Iterative CTF refinement and Bayesian polishing were performed for each state until the resolution converged. To prevent overfitting, SIDESPLITTER [49] was integrated into all subsequent 3D refinements for the closed and intermediate states. Particles were unbinned during Bayesian polishing to a pixel size of 0.9 Å/px. This resulted in a final resolution of 2.6 Å for the closed state from 79,604 particles and 2.8 Å for the intermediate from 113,615 particles, following the gold standard Fourier shell correlation cut-off at 0.143. Local resolution was calculated in Relion.

### Model building and validation

For model building of both the closed and intermediate states, a previously published SthK structure (PDB: 7tj5) was docked into the final cryo-EM density maps using ChimeraX [50] and manually adjusted in COOT [51]. Residues that could not be resolved in previous structures were added, and the resulting coordinates were real-space refined in PHENIX [52, 53]. cAMP and lipids were manually added to each model in COOT, and outliers were fixed. The final models were validated using MolProbity [54]. Pore diagrams were made using HOLE and Cα RMSD calculations were performed using ChimeraX [50, 55]. All structural figures were prepared using ChimeraX.

## Acknowledgements

We thank C. Fluck (Weill Cornell Medicine, Cryo-EM Core Facility) for advice on cryo-EM sample preparation and screening, Cryo-EM data collection at NYU Langone Health’s Cryo-Electron Microscopy Laboratory (RRID: SCR-019202), W. Zagotta and E. Evans (University of Washington, Department of Physiology and Biophysics) for providing us with *E. coli* C43 cyA^−^ cells, and R. Rusinova (Weill Cornell Medicine, Department of Physiology and Biophysics) for guidance on proteoliposome preparation and DLS measurements. This work was sponsored by the NIH (grant no. GM124451 to C.M.N.) and the National Science and Technology Council of Taiwan (formerly Ministry of Science and Technology, grant no. 111-2917-I-564-001 to C.-C.L).

## Author contributions

C.-C.L. and C.M.N designed the research and wrote the manuscript. C.-C.L. performed the experiments and analyzed the data.

---

**Extended Data Figure 1.**
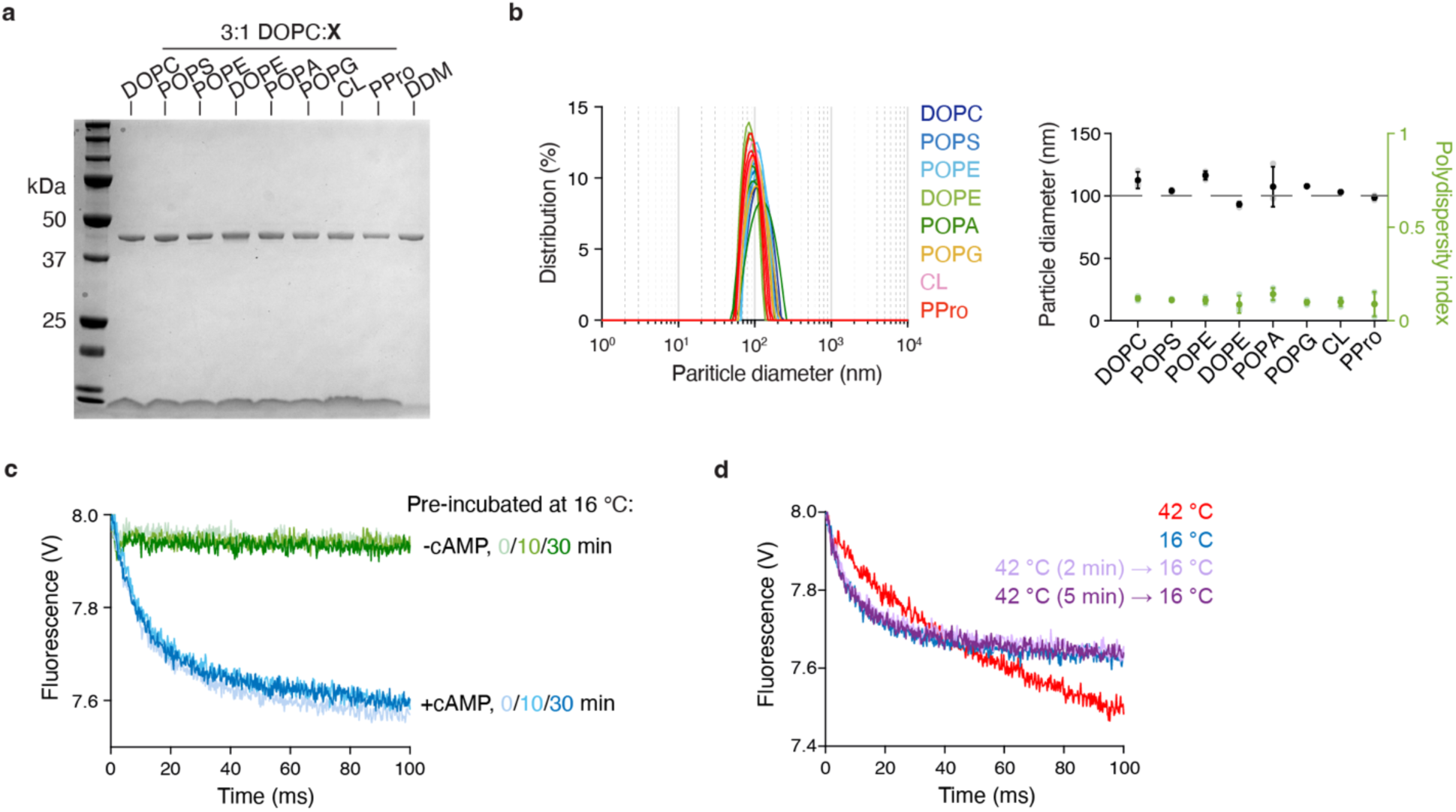
Properties of SthK proteoliposomes for functional assay. **a**, SDS-PAGE analysis of SthK reconstituted into various lipid compositions (PC-only or 3:1 DOPC:X, with X = POPS, POPE, DOPE, POPA, POPG, CL, or PPro) for the stopped-flow Tl^+^ flux assay. Samples were prepared as described for the stopped-flow assay, except that reconstitution buffer was added instead of ANTS solution. After extrusion, protein was extracted by adding 30 mM DDM and incubated for 1.5 hours under constant agitation. Samples were cleared through a Spin-X column and analyzed by SDS-PAGE to assess protein incorporation across lipid conditions. Purified SthK in DDM was included as reference. **b**, Left, proteoliposome size determined using dynamic light scattering. Proteoliposomes were prepared as above and diluted to ∼30 μg/ml lipid after extrusion. The experiment was independently repeated three times, with individual experiments shown on the left. The average size and polydispersity index, along with their standard deviations (s.d.), are displayed on the right. **c**, Cold-desensitization test of SthK in 3:1 DOPC:POPE. Quenching kinetics from stopped-flow flux assays for SthK WT pre-incubated at 16 °C with (blue) or without (green) 200 μM cAMP for indicated durations, then assayed at 16 °C. Experiments were repeated three times independently; a representative result is shown. **d**, Reversibility test of SthK. Quenching kinetics from stopped-flow flux assays for SthK WT assayed directly at 16 °C (blue), at 42 °C (red), or after pre-incubation at 42 °C for 2 min (pink) or 5 min (purple) followed by assay at 16 °C, all performed in the presence of 200 µM cAMP. Experiments were repeated three times independently; a representative result is shown.

**Extended Data Figure 2.**
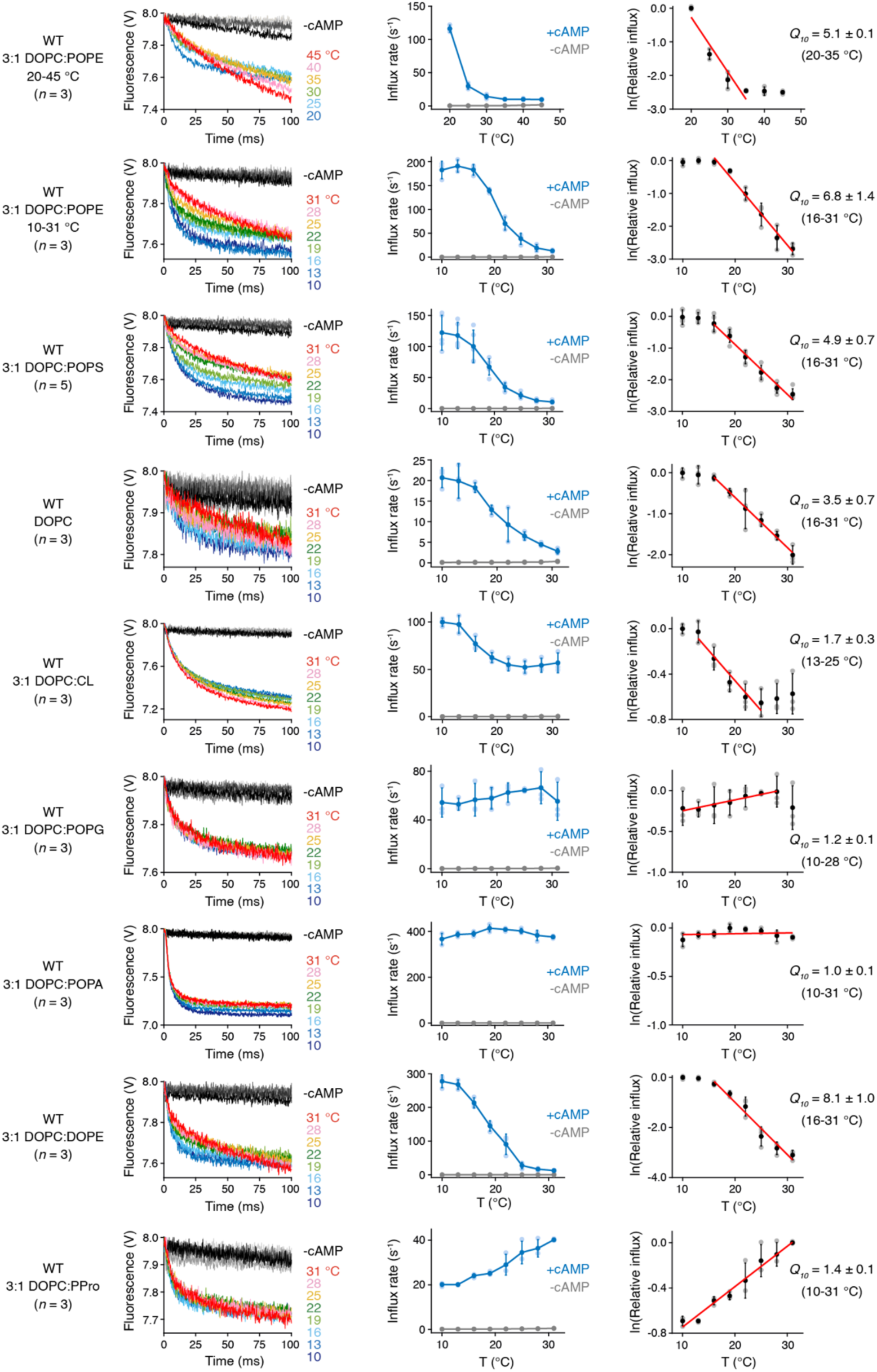
Stopped-flow functional assays of SthK WT. Left: Representative quenching kinetics from stopped-flow flux assays for SthK WT in different lipid compositions. Results obtained at different temperatures and activated by 200 μM cAMP are shown in multiple colors. Traces without cAMP are depicted in grayscale, with darker shades indicating higher temperatures. Center: Corresponding initial Tl⁺ flux rates. Filled symbols represent mean ± s.d. for biological replicates (*n* = 3 for all except PS, where *n* = 5). Right: Logarithmic plots of relative influx rates at various temperatures, with fitting curves used to determine the temperature coefficient (*Q_10_*) and the corresponding fitting range. *Q_10_* values are shown as mean ± s.d.

**Extended Data Figure 3.**
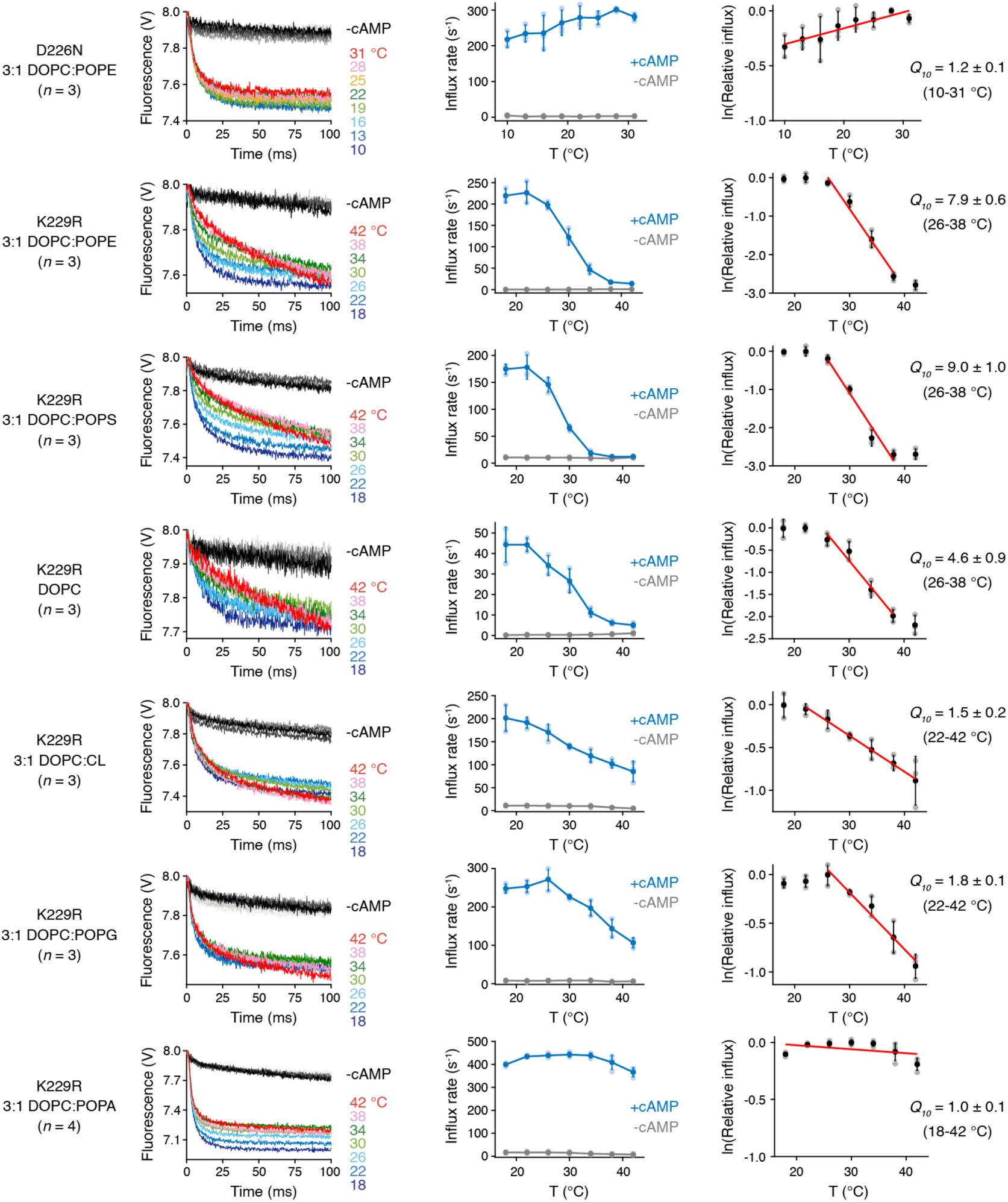
Stopped-flow functional assays of SthK variants. Left: Representative quenching kinetics from stopped-flow flux assays for SthK variants in different lipid compositions. Results obtained at different temperatures and activated by 200 μM cAMP are shown in multiple colors. Traces without cAMP are depicted in grayscale, with darker shades indicating higher temperatures. Center: Corresponding initial Tl⁺ flux rates. Filled symbols represent mean ± s.d. for biological replicates (*n* = 3 for all except K229R in PA, where *n* = 4). Right: Logarithmic plots of relative influx rates at various temperatures, with fitting curves used to determine the temperature coefficient (*Q_10_*) and the corresponding fitting range. *Q_10_* values are shown as mean ± s.d.

**Extended Data Figure 4.**
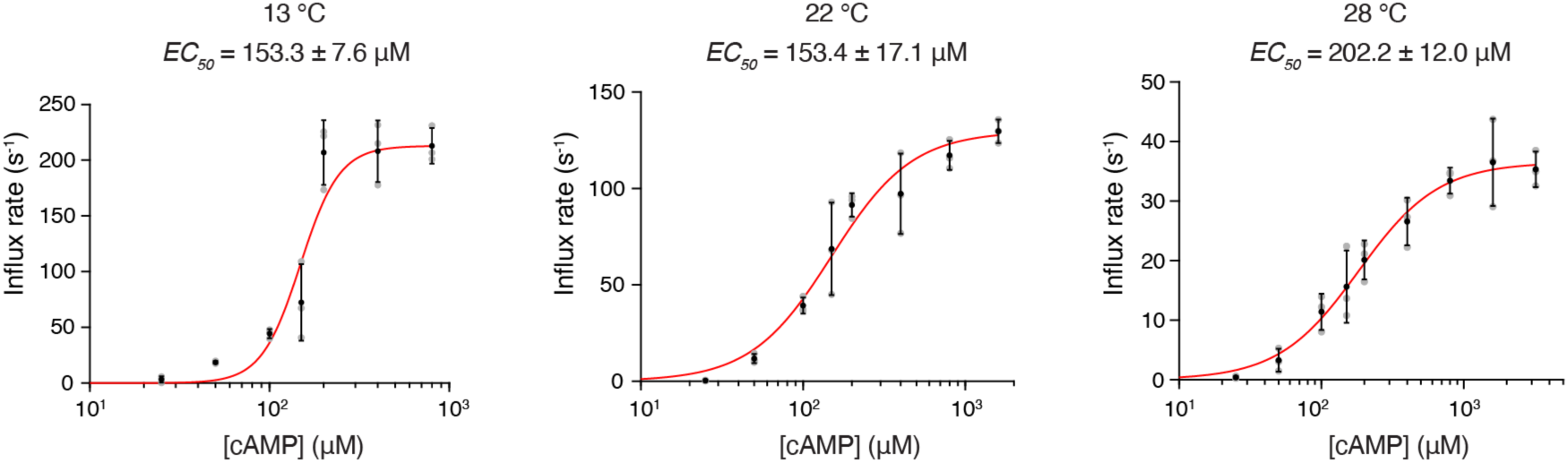
*EC50* for cAMP activation has little temperature dependence. Initial Tl^+^ flux rates of SthK were measured in liposomes composed of DOPC:POPE at a 3:1 ratio across a range of cAMP concentrations. Each experiment was repeated independently three times, and *EC₅₀* values were obtained by fitting individual replicates using the Hill equation, reported values represent mean ± s.d. across replicates. The Hill coefficients at 22 °C and 28 °C were 1.8 ± 0.3 and 1.5 ± 0.3, respectively. At 13 °C, the Hill coefficient was constrained to 4, reflecting the maximal number of cAMP binding sites, as unconstrained fitting produced unrealistically high values (>10). A two-tailed unpaired t-test revealed no significant difference in *EC₅₀* between 13 °C and 22 °C (*p* = 0.996, *t* = –0.005).

**Extended Data Figure 5.**
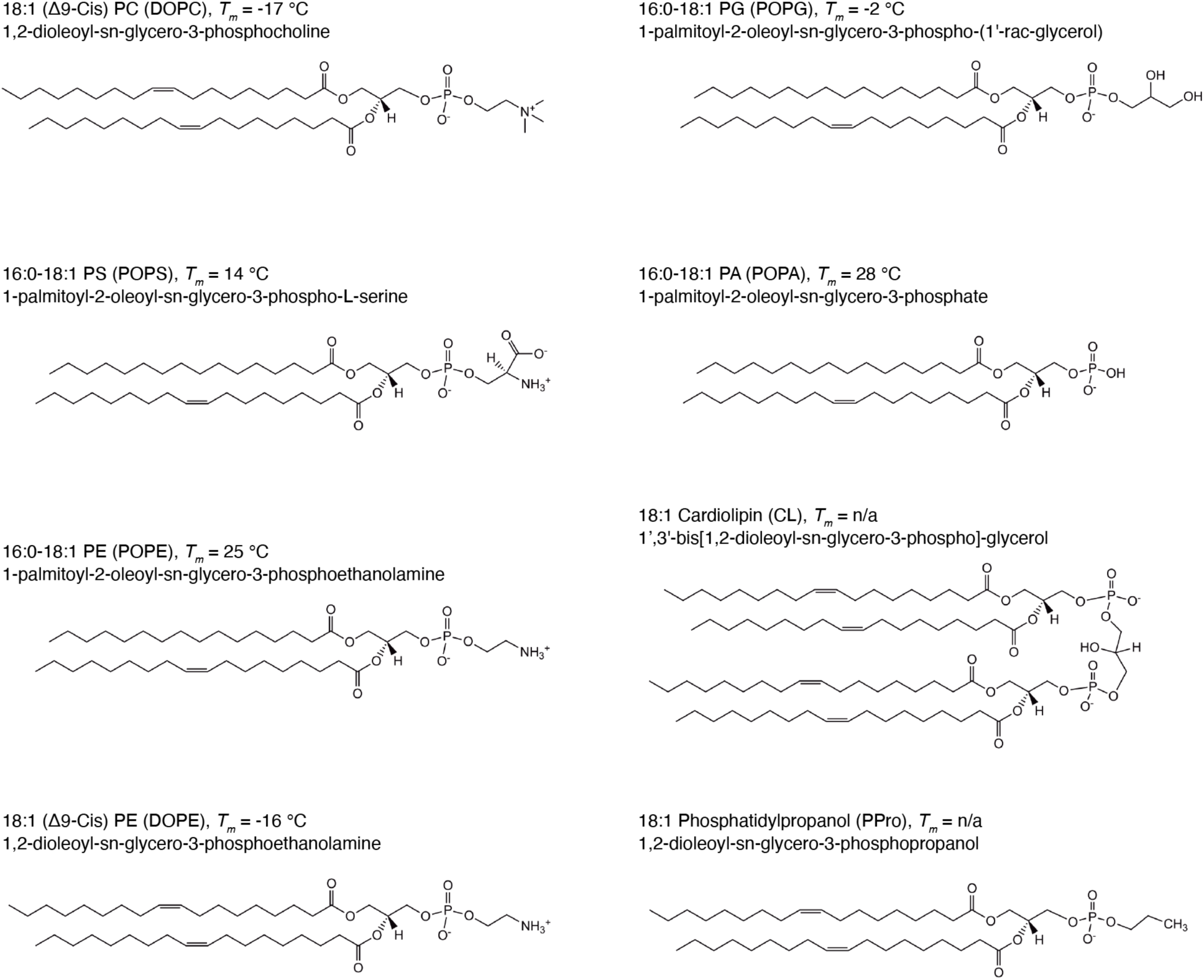
Characteristics of different lipids. Chemical structures of lipids utilized in this study, along with their IUPAC and simplified abbreviations, and phase transition temperatures (*T_m_*). Information was obtained from: https://avantilipids.com.

**Extended Data Figure 6.**
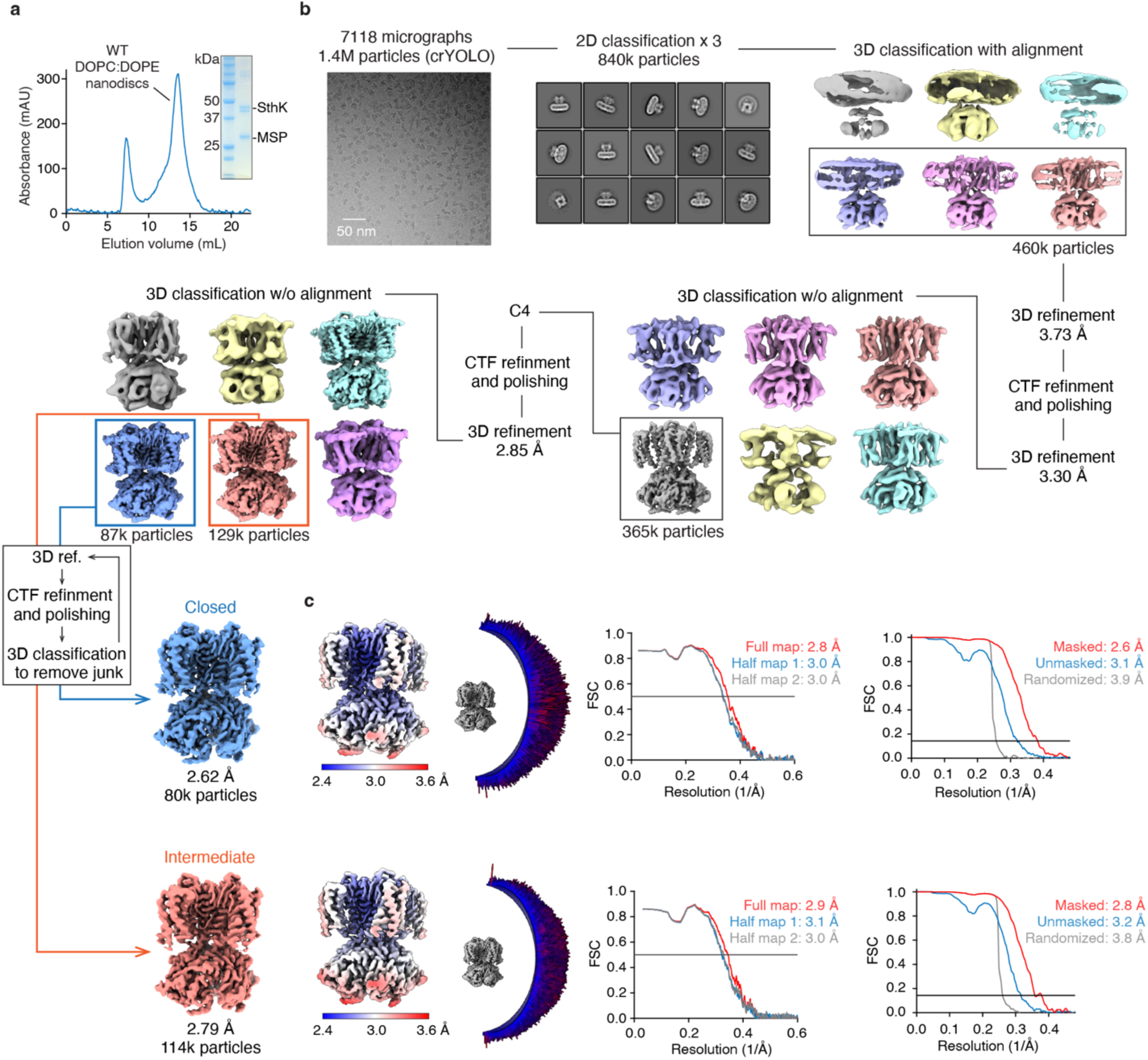
Cryo-EM data processing workflow of cAMP-bound SthK WT in the presence of PE lipids. **a**, Size-exclusion chromatography profile and SDS-PAGE of purified SthK WT in 1:1 DOPC:DOPE nanodiscs. **b**, The cryo-EM processing workflow of SthK WT in 1:1 DOPC:DOPE nanodiscs in the presence of 3 mM cAMP. **c**, From left to right: local resolution maps, angular distributions of the final maps, FSC curves between the model and the final map (full map), as well as the two half maps (half map 1 and half map 2), and FSC curves of the final maps for the closed (top) and intermediate (bottom) states.

**Extended Data Figure 7.**
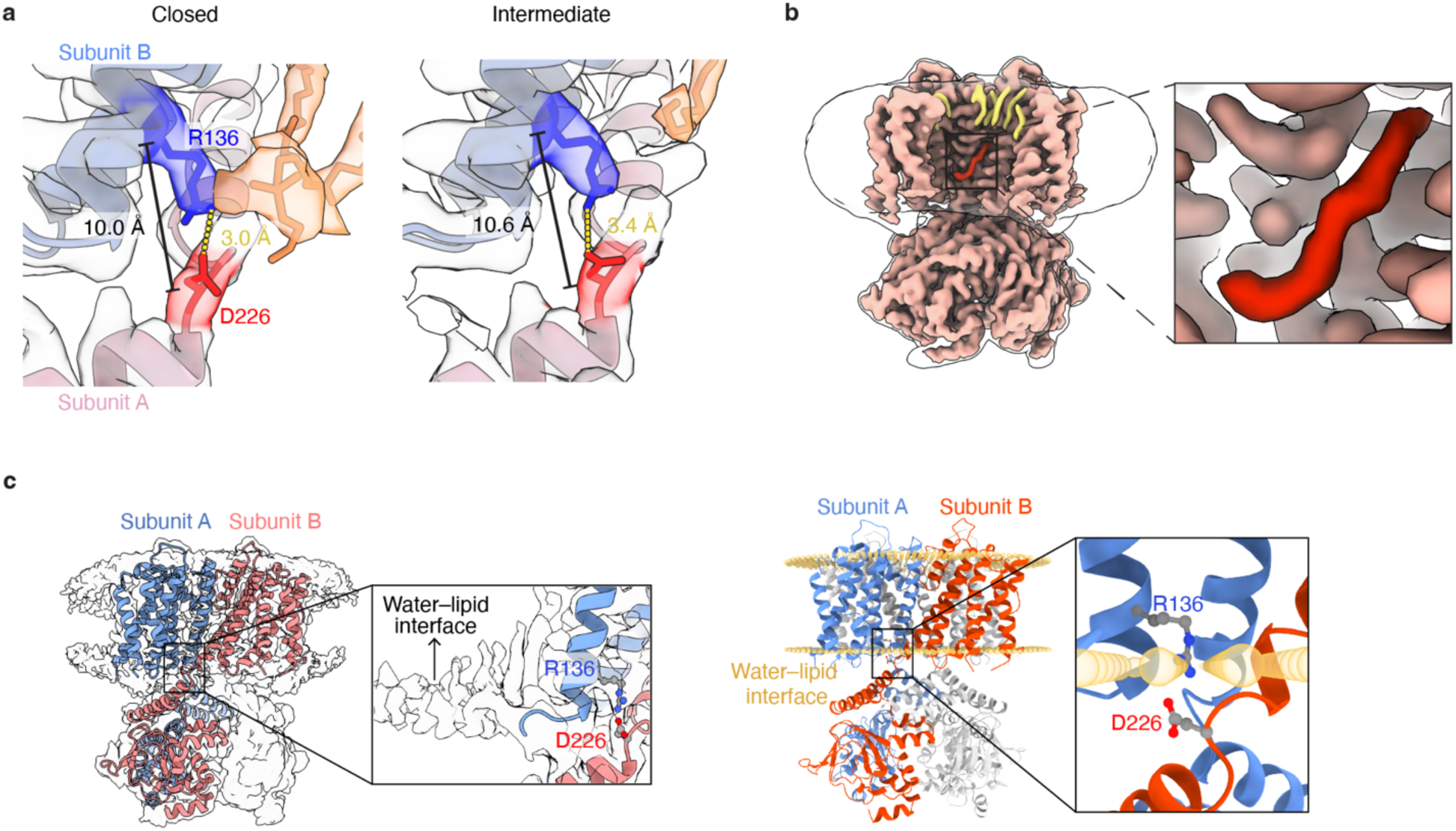
Features of SthK structures in the presence of PE lipids. **a**, Salt bridge distance is shorter in the closed state compared to intermediate state. Cryo-EM density maps and structural models of the intersubunit salt bridge R136–D226 in the closed (left) and intermediate (right) states. R136 and D226 are highlighted in blue and red, respectively; surrounding lipids are shown in orange. Side-chain and Cα–Cα distances are labeled. **b**, There is also a lipid density near the salt bridge in the intermediate state. Protein density is shown in pink. Most lipid densities are shown in yellow, with the lipid density in the inner leaflet near the intersubunit salt bridge R136–D226 highlighted in red. The inset is a zoom-in highlighting this red lipid density. **c**, Left: Cryo-EM density of SthK in PE-containing nanodiscs at a lower density threshold, highlighting in the zoomed-in inset the proximity of the R136–D226 salt bridge to the water–lipid interface. For clarity, only two adjacent subunits of the SthK model are shown in red and blue cartoon representation. Right: Prediction of SthK (PDB: 7tj5) embedded in a DOPC bilayer using the Orientations of Proteins in Membranes (OPM) database. Water–lipid interfaces are shown as yellow spheres. The zoomed-in view in inset shows the intersubunit salt bridge R136–D226 is also located at water–lipid interface in this prediction.

**Extended Data Figure 8.**
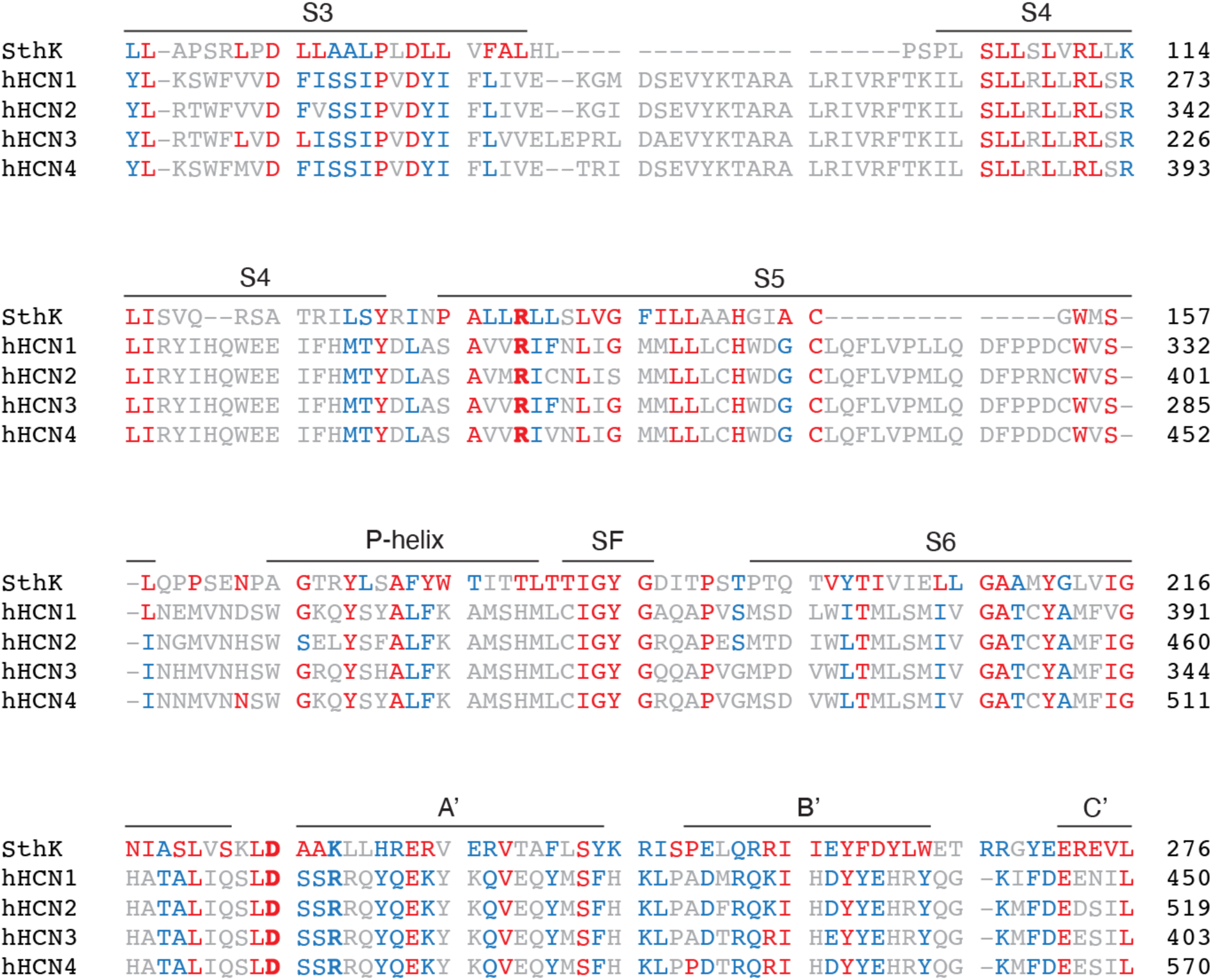
Sequence homology between SthK and eukaryotic channels. Sequence alignment of SthK and human CNG and HCN channels based on UniProt entries: SthK (G0GA88), human HCN1 (O60741), human HCN2 (Q9UL51), human HCN3 (Q9P1Z3), and human HCN4 (Q9Y3Q4). The alignment was performed using CLUSTAL O (1.2.4). Identical residues are colored in red, similar residues in blue. The residues forming intersubunit salt bridges in SthK and their conservation are highlighted in bold.

**Extended Data Table 1.**
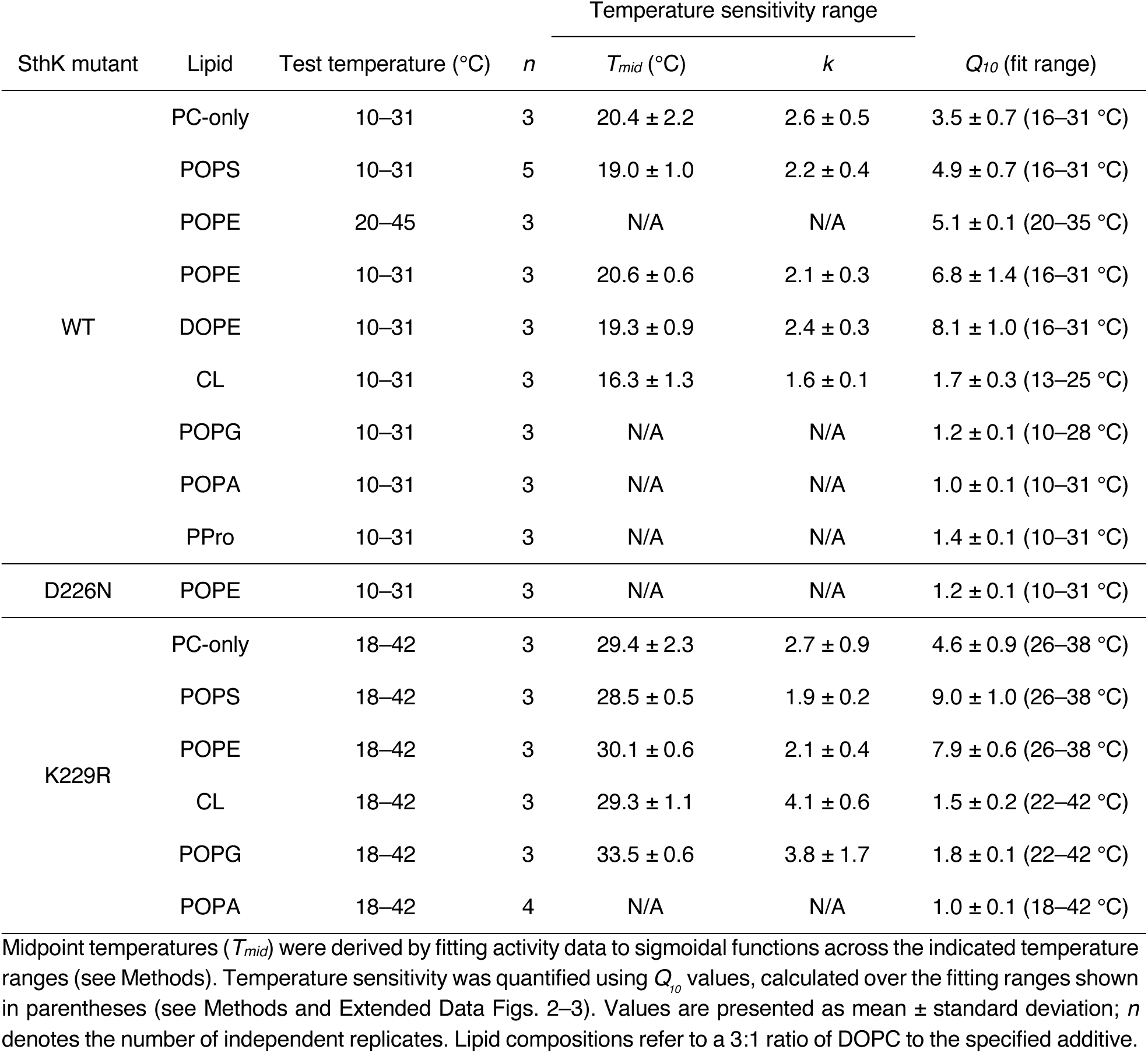
Summary of the temperature-sensitive range and temperature sensitivity of SthK in different lipid environments.

| SthK mutant | Lipid | Test temperature (°C) | *n* | Temperature sensitivity range |  |  |
| --- | --- | --- | --- | --- | --- | --- |
| | | | | $T_{mid}$ (°C) | *k* | $Q_{10}$ (fit range) |
| WT | PC-only | 10–31 | 3 | 20.4 ± 2.2 | 2.6 ± 0.5 | 3.5 ± 0.7 (16–31 °C) |
|  | POPS | 10–31 | 5 | 19.0 ± 1.0 | 2.2 ± 0.4 | 4.9 ± 0.7 (16–31 °C) |
|  | POPE | 20–45 | 3 | N/A | N/A | 5.1 ± 0.1 (20–35 °C) |
|  | POPE | 10–31 | 3 | 20.6 ± 0.6 | 2.1 ± 0.3 | 6.8 ± 1.4 (16–31 °C) |
|  | DOPE | 10–31 | 3 | 19.3 ± 0.9 | 2.4 ± 0.3 | 8.1 ± 1.0 (16–31 °C) |
|  | CL | 10–31 | 3 | 16.3 ± 1.3 | 1.6 ± 0.1 | 1.7 ± 0.3 (13–25 °C) |
|  | POPG | 10–31 | 3 | N/A | N/A | 1.2 ± 0.1 (10–28 °C) |
|  | POPA | 10–31 | 3 | N/A | N/A | 1.0 ± 0.1 (10–31 °C) |
|  | PPro | 10–31 | 3 | N/A | N/A | 1.4 ± 0.1 (10–31 °C) |
| D226N | POPE | 10–31 | 3 | N/A | N/A | 1.2 ± 0.1 (10–31 °C) |
| K229R | PC-only | 18–42 | 3 | 29.4 ± 2.3 | 2.7 ± 0.9 | 4.6 ± 0.9 (26–38 °C) |
|  | POPS | 18–42 | 3 | 28.5 ± 0.5 | 1.9 ± 0.2 | 9.0 ± 1.0 (26–38 °C) |
|  | POPE | 18–42 | 3 | 30.1 ± 0.6 | 2.1 ± 0.4 | 7.9 ± 0.6 (26–38 °C) |
|  | CL | 18–42 | 3 | 29.3 ± 1.1 | 4.1 ± 0.6 | 1.5 ± 0.2 (22–42 °C) |
|  | POPG | 18–42 | 3 | 33.5 ± 0.6 | 3.8 ± 1.7 | 1.8 ± 0.1 (22–42 °C) |
|  | POPA | 18–42 | 4 | N/A | N/A | 1.0 ± 0.1 (18–42 °C) |

**Extended Data Table 2.**
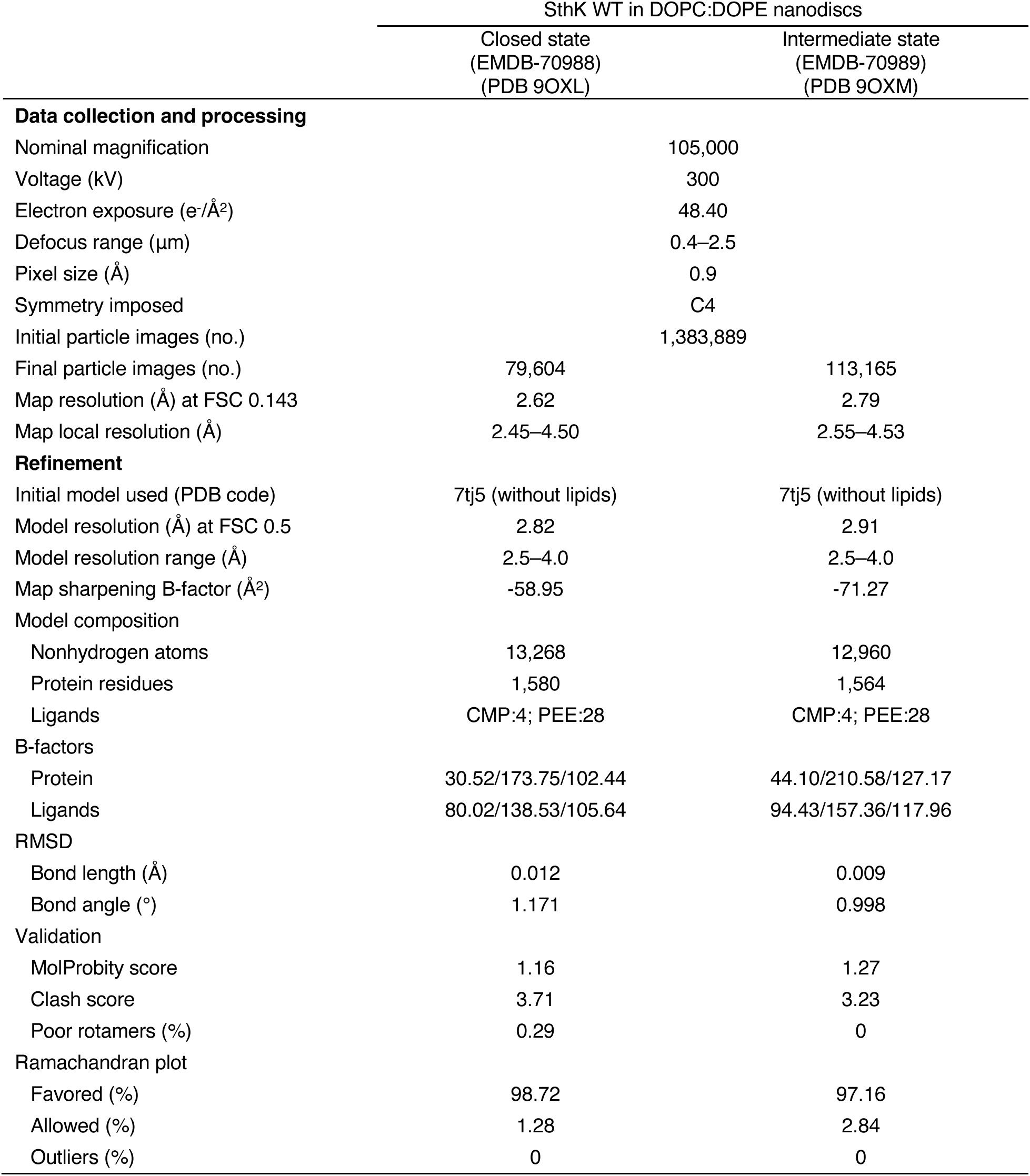
Cryo-EM data collection, refinement and validation statistics.

